# A mutation-agnostic and allele-specific ASO strategy demonstrates potent functional rescue and retinal preservation in RHO-linked retinitis pigmentosa

**DOI:** 10.64898/2026.08.25.747013

**Authors:** Salome Spaag, Wen-Hsuan Wu, Justin Yun, Aykut Demirkol, Thomas Winogrodzki, Anders Steen Knudsen, Georgy Komissarov, Chyuan-Sheng Lin, Melita Kaltak, Maddalena Fuso, Kaushambee Dave, Katarina Stingl, Angela Armento, Laura Kühlewein, Britta Baumann, Carmen Ayuso, Lidia Fernandez-Caballero, Rob Collin, Femke Bukkems, Susanne Roosing, Zelia Corradi, Sandro Banfi, Francesca Simonelli, Clemens Lochmann, Marianthi Karali, Sylvia Bolz, Susanne Kohl, Eberhart Zrenner, Bernd Wissinger, Kevin Achberger, Stephen H. Tsang, Pietro De Angeli

**Affiliations:** Institute for Ophthalmic Research, Centre for Ophthalmology, University Hospital Tübingen, 72076 Tübingen, Germany; Department of Biomedical Engineering, Columbia University, New York, New York, USA; Jonas Children’s Vision Care and Bernard & Shirlee Brown Glaucoma Laboratory, Institute of Human Nutrition, Columbia Stem Cell Initiative, New York, New York, USA; Edward S. Harkness Eye Institute, Columbia University Irving Medical Center, New York-Presbyterian Hospital, New York, New York, USA; Departments of Ophthalmology, Pathology & Cell Biology, Vagelos College of Physicians and Surgeons, Columbia University Irving Medical Center, New York, New York, USA; University Eye Hospital, Center for Ophthalmology, University of Tuebingen, Germany; Institute of Neuroanatomy & Developmental Biology (INDB), Eberhard Karls University Tübingen, Tübingen, Germany; Division Translational Genomics of Neurodegenerative Diseases, Hertie-Institute for Clinical Brain Research and Center of Neurology, University of Tübingen, Tübingen, Germany; Department of Genetics & Genomics, Instituto de Investigación Sanitaria-Fundación Jiménez Díaz University Hospital, Universidad Autónoma de Madrid (IIS-FJD, UAM), Madrid, Spain; Astherna B.V., Nijmegen, the Netherlands; Department of Human Genetics, Radboud University Medical Center, 6525 GA, Nijmegen, the Netherlands; Department of Precision Medicine, University of Campania "Luigi Vanvitelli", Naples, Italy; Telethon Institute of Genetics and Medicine, Pozzuoli, Italy; Center for Biomedical Network Research on Rare Diseases (CIBERER), Instituto de Salud Carlos III, Madrid, Spain.; Eye Clinic, Multidisciplinary Department of Medical, Surgical and Dental Sciences, University of Campania “Luigi Vanvitelli”, Naples, Italy

## Abstract

Autosomal dominant retinitis pigmentosa (adRP) caused by *RHO* mutations is a leading form of inherited retinal degeneration. Extensive allelic heterogeneity of *RHO* pathogenic variants limits the translational applicability of mutation-specific gene therapies. To address this, we developed SNARE (SNP-guided Silencing of Aberrant RHO Expression), a mutation-independent, allele-specific antisense oligonucleotide (ASO) strategy. SNARE selectively suppresses mutant *RHO* transcripts by targeting the common, benign c.-26A/G single-nucleotide polymorphism (SNP) as an allelic discriminator. Candidate gapmer ASOs were screened in engineered reporter lines and validated in patient-derived retinal organoids, identifying RHOligo-A as the lead c.-26A-targeting candidate. In *vitro*, RHOligo-A achieved robust, preferential knockdown of the target allele, improving RHO localization in retinal organoids, and demonstrated a favorable safety profile with minimal transcriptomic off-target effects and no detectable immunostimulatory activity. Subsequent validation in a novel, humanized RHO^P347L/WT^ mouse model, achieved sustained c.-26A-linked allele-selective suppression, retinal structure preservation, and significantly restored visual function, upon a single intravitreal administration. These findings establish RHOligo-A and SNARE as a scalable, mutation-independent therapeutic platform with strong translational potential and substantial clinical reach for RHO-associated adRP.

## INTRODUCTION

Retinitis pigmentosa (RP) is the most prevalent form of inherited retinal diseases (IRD), affecting approximately 1 in 3,000–4,000 individuals worldwide and accounting for roughly 25– 35% of all IRD patients(1–3). Pathogenic variants in more than 100 genes have been implicated in RP, which can be inherited as an autosomal dominant, autosomal recessive, or X-linked trait (2, 4, 5). Autosomal dominant RP (adRP) constitutes about 20% of all RP cases, among which mutations in the *RHO* gene (OMIM*180380, #613731) represent approximately 20–30%, corresponding to a general population prevalence of ∼1 in 80,000–100,000 individuals (6–10). Most pathogenic *RHO* variants are missense substitutions that act through dominant-negative or gain-of-function mechanisms, whereby mutant RHO interferes also with wild-type protein function. Molecular disease mechanisms include protein misfolding and endoplasmic reticulum retention, defective trafficking to the photoreceptor outer segment, and abnormal constitutive activation of the phototransduction cascade (11, 12). Because of this dominant-negative behavior, gene augmentation is not a suitable therapeutic strategy. In contrast, reduced RHO dosage is tolerated, as heterozygous loss-of-function variants and early nonsense mutations do not cause IRD (13, 14).

Consequently, therapeutic development has aimed at selectively reducing mutant *RHO* expression, including allele-specific gene editing and RNA-mediated silencing strategies such as antisense oligonucleotides (ASOs). However, the mutational spectrum of *RHO* is highly heterogeneous, with more than 250 pathogenic variants described, and mutation-specific approaches therefore addresses primarily a small number of recurrent variants such as P23H (15–17). This mutation-by-mutation paradigm limits scalability and patient reach, as the small number of individuals per variant reduces the feasibility of advancing each candidate beyond proof-of-concept studies.

To overcome variant heterogeneity, mutation-independent “ablate-and-replace” strategies have been proposed, in which both endogenous *RHO* alleles are disrupted or knocked down while a suppression-resistant wild-type *RHO* cDNA is delivered, typically via adeno-associated viral (AAV) vectors (18, 19). Although theoretically genotype-agnostic, these approaches face important practical limitations. Precise control of AAV-driven RHO expression is difficult to achieve, and even overexpression can be toxic to photoreceptors, as demonstrated in animal models and by the presence of adRP patients carrying four copies of wild-type *RHO* (20). In addition, delivery is commonly performed by subretinal injection, restricting treatment and later visual field recovery to a limited retinal area (21).

An alternative mutation-independent strategy, which might be able to target a substantial proportion of *RHO*-adRP patients, is enabled by the presence of benign heterozygous single-nucleotide polymorphisms (SNPs) within the gene, which can serve as allele-discriminating “entry points” when pathogenic variants are in *cis* with a given SNP allele (22, 23). Gapmer ASOs, which recruit RNase H to induce degradation of target RNA, are particularly well suited for single-nucleotide discrimination and selective transcript knockdown (15, 22, 24). Importantly, multiple ASO-based therapeutics have already demonstrated safety and clinical feasibility for ocular indications in human trials, supporting this modality as a translatable therapeutic platform (25–28). Leveraging these two aspects, we developed **SNARE (SNP-guided silencing of Aberrant RHO Expression)**, a mutation-independent ASO strategy designed to selectively suppress mutant *RHO* transcripts while sparing the healthy allele. The high prevalence of the target heterozygous SNP enables broad patient applicability, positioning SNARE as a scalable therapeutic strategy for *RHO*-associated adRP. We demonstrate potent and allele-selective activity across multiple *in vitro* models and translate these findings *in vivo* using a fully humanized RHO:p.P347L mouse model, where treatment, using the lead RHOligo-A ASO candidate, resulted in functional rescue and preservation of retinal structure.

## RESULTS

### Determination of frequent SNPs in the *RHO* gene as a basis for mutation-agnostic genetic therapeutics

To identify common *RHO* SNPs for allele discrimination, we queried gnomAD v4.1.0 and identified five benign minor alleles with heterozygous frequencies between 2.71% and 21.65%. To confirm allele frequency in patients, we screened these SNPs in a clinically relevant cohort of 20 unrelated RHO-adRP patients from the local Tübingen RetDis database. The SNPs c.-26A/G (rs7984), c.696+4C/T (rs56340615), c.-51G/A (rs2269736), c.360C>T (p.Gly120Gly, rs79765751), and c.923-23G/A (rs2071092) were detected in 8, 3, 1, 1 and 0 patients, respectively.

Because c.-26A/G was the most frequent SNP, we expanded genotyping to a total of 445 patients across four independent cohorts: Tübingen (n=205), Madrid (n=81), Nijmegen (n=106), and Naples (n=53). Heterozygosity for c.-26A/G was present in 25.3% (n=52), 26.6% (n=21), 38.7% (n=41), and 24.5% (n=13) of patients, respectively, yielding a cumulative frequency of 28.5% (n=127; **Figure 1A**). Across these cohorts, we identified 66 unique pathogenic *RHO* variants. The most prevalent were c.1040C>T (p.P347L; n=21, 4.7%) and c.403C>T (p.R135W; n=11, 2.5%); all others were private or rare.

**Figure 1.**
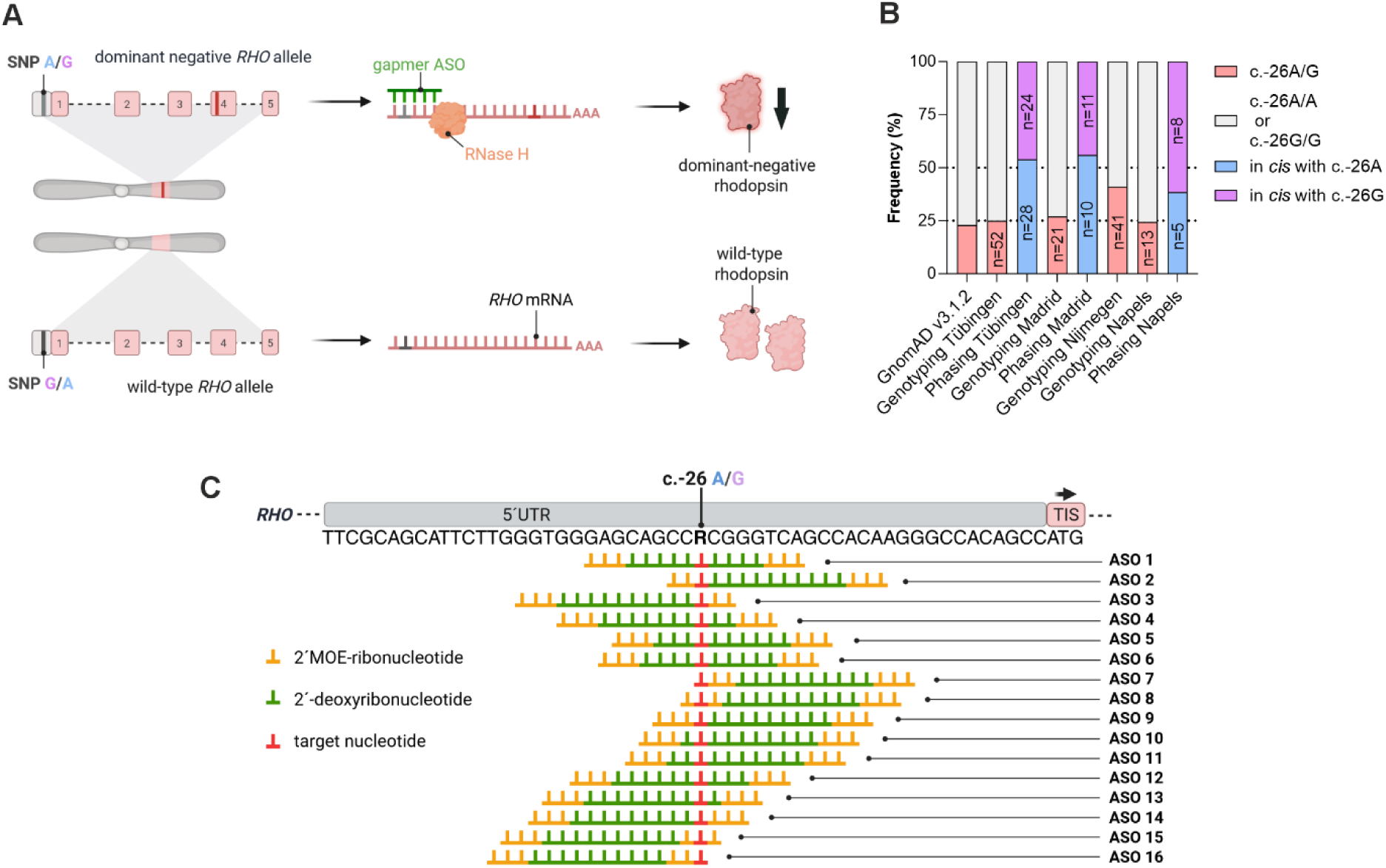
Outline of SNP-guided Silencing of Aberrant Expression (SNARE). (A) Heterozygous frequency of the targeted single nucleotide polymorphism (SNP; rs7984) in four different *RHO* patient population compared with allele frequency data from gnomAD v3.1.2. For three patient cohorts, phasing between rs7984 and pathogenic *RHO* variants is also shown. (B) Schematic representation of the SNARE molecular mechanism. Allele-specific antisense oligonucleotides selectively target either c.-26A or c.-26G *RHO* transcripts, depending on which allele carries the pathogenic *RHO* variant. Selective knockdown of the mutant transcript results in reduction of the mutant RHO protein. (C) Design of allele-specific ASOs targeting either the c.-26A or c.-26G allele, with their corresponding complementary binding sequences located in the 5′ UTR of the *RHO* transcript. TIS = translation initiation site.

Subsequent phasing analysis showed the mutant *RHO* allele was in *cis* with the c.-26A allele in 54% (n=28) of Tübingen, 56% (n=10) of Madrid, and 39% (n=5) of Naples patients (**Figure 1A**). Phasing was not available for the Nijmegen cohort.

### Design and *in vitro* screening of *RHO*:c.-26A/G-targeting gapmer ASOs

To maximize single-nucleotide discrimination at c.-26A/G, we designed a 1-bp walk of 16-mer gapmer ASOs, each containing a 10-mer DNA gap flanked by 3-mer modified RNA wings to support RNase H1 recruitment, nuclease stability, and target affinity (29–31). This design generated 32 candidates (16 per allele; **Supplementary Table 1**) that were tested for knockdown and allele-specificity (**Figure 1B–C**).

Because *RHO* expression is restricted to retinal photoreceptors, we established two stable HEK293T reporter cell lines (32). These lines express full-length *RHO* alleles—either *RHO*:c.[- 26A;696+4T] (A line) or *RHO*:c.[-26G;696+4C] (G line)—fused at the 3′ end to mCherry via a flexible linker. Transgenes were integrated using reverse-complement lentiviral vectors to ensure intron retention during production, alongside a zeocin resistance gene for clonal selection (**Figure 2A**). Selected single-cell clones were validated for *RHO* expression via qRT- PCR, for full-length transgene integration and copy numbers, correct splicing, and RHO- mCherry co-localization (**Figure 2B–F**). These reporter lines were used as controlled, high- throughput allele-discrimination tools; endogenous validation was therefore performed subsequently in patient-derived retinal organoids and a humanized *in vivo* model.

**Figure 2.**
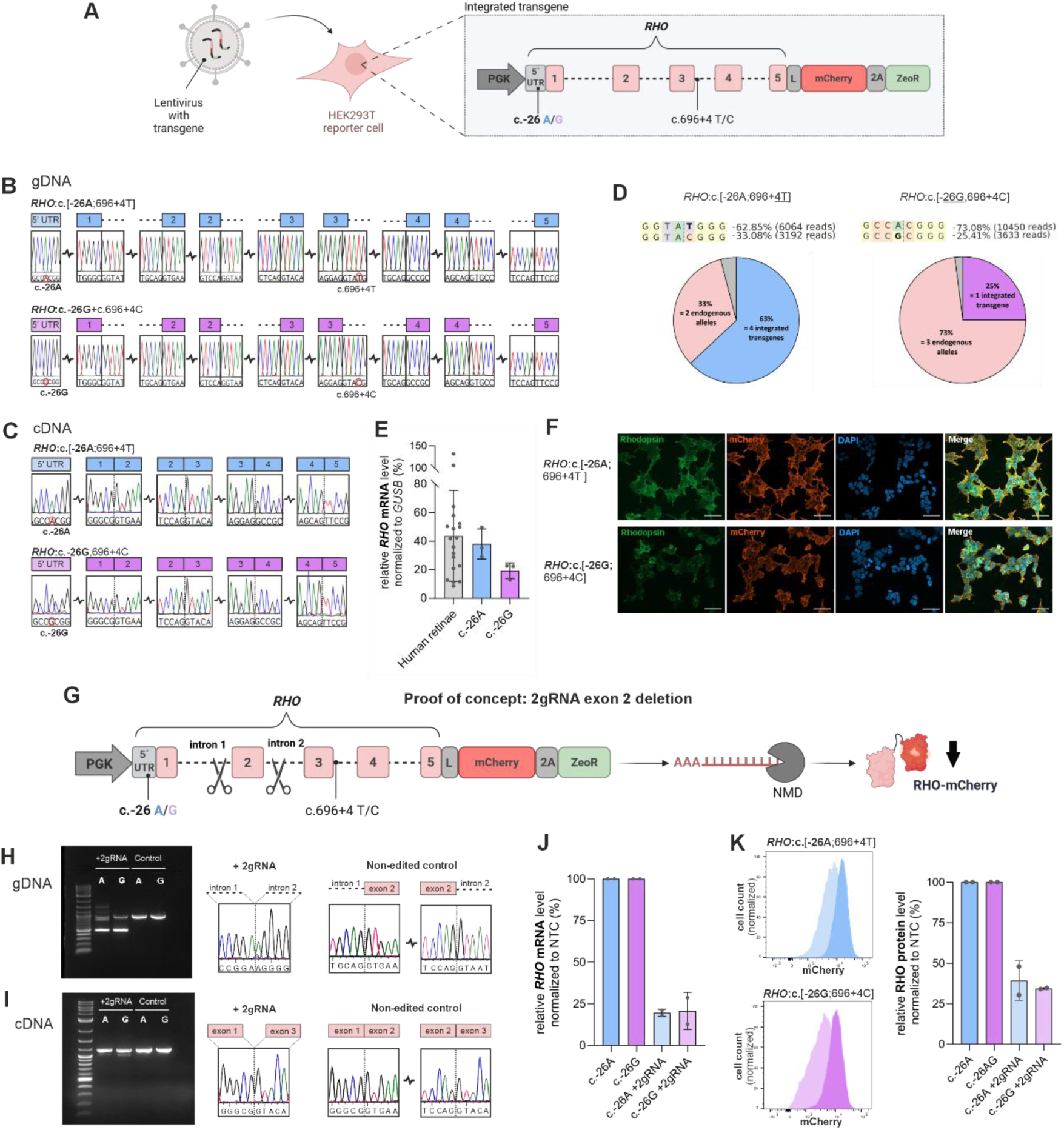
Generation and characterization of the *RHO* reporter cell lines. (A) Schematic representation of the lentiviral constructs used to generate the reporter lines *RHO*:c.[-26A;696+4T] and *RHO*:c.[-26G;696+4C]. The full-length *RHO* gene, including the 5′ untranslated region (UTR) and intronic sequences, was fused via a flexible linker (L) to the fluorescent protein mCherry. A self-cleaving 2A peptide sequence downstream of mCherry enables co-expression of the antibiotic resistance gene ZeoR. (B) Genomic sequencing of the RHO–mCherry transgene in established reporter lines, highlighting intron–exon boundaries and the c.-26 position in the 5′ UTR. (C) cDNA sequencing of *RHO*–*mCherry* transcripts, indicating exon–exon junctions and the c.-26 position in the 5′ UTR. (D) Transgene-to-endogenous *RHO* gene ratio determined by high-throughput sequencing. Ratios were calculated from sequencing reads of the heterozygous single-nucleotide polymorphisms rs56340615 c.696+4C/T and c.- rs7984 26A/G in the *RHO*:c.[-26A;696+4T] and *RHO*:c.[-26G;696+4C] lines, respectively. (E) Expression levels of the *RHO–mCherry* transgene in reporter cell lines compared with human retina. (F) Immunofluorescence staining of RHO reporter cell lines showing RHO and mCherry protein expression and co-localization. (G) Schematic of the dual-gRNA/Cas9 strategy used to validate reporter responsiveness. Scissors indicate gRNA target sites. Deletion of exon 2 induces a frameshift and premature stop codons, triggering nonsense-mediated mRNA decay (NMD) and reduced RHO–mCherry protein levels. (H–K) Validation of reporter lines following genome editing: (H) genomic DNA analysis and (I) transcript analysis by agarose gel electrophoresis and fragment sequencing; (J) quantification of transgene transcripts by qRT–PCR; and (K) RHO–mCherry protein levels assessed by flow cytometry. For panels (E,J,K), Data are presented as mean ± standard deviation. Individual dots represent individual measurement.

To confirm reporter responsiveness, we used CRISPR/Cas9 to excise exon 2, inducing a frameshift (**Figure 2G–H**). Agarose gel electrophoresis confirmed DNA-level deletion, while quantitative RT-PCR revealed substantial nonsense-mediated mRNA decay (33), leaving only 9.5 ± 1.5% (A line) and 20.7 ± 7.9% (G line) residual transcript (**Figure 2I–J**). Correspondingly, flow cytometry demonstrated a robust reduction in mCherry fluorescence to residual levels of 39 ± 12% (A line) and 34 ± 1% (G line), validating the platform (**Figure 2K**).

For ASO screening, reporter cells were transfected with 50 nM ASO in serum-deprived medium to limit proliferation dilution, and analyzed 72 h post-transfection. In initial protein- level potency screens by flow cytometry, candidate c.-26A ASOs (ASO_5, _9, _11, _15) achieved target-allele knockdowns of 40.5 ± 5%, 31.1 ± 8.7%, 42.6 ± 4.9%, and 35.8 ± 5.9%, respectively (**Figure 3A, Supplementary Figure 1**). Candidate c.-26G ASOs (ASO_2, _3, _5, _9, _11, _13, _15) yielded knockdowns of 50 ± 6.7%, 52.7 ± 5.6%, 64 ± 11%, 49.2 ± 14.7%, 49.3 ± 24.3%, 46.2 ± 6.7%, and 65.5 ± 1.7%, respectively (**Figure 3B**, **Supplementary Figure 2**). When evaluated for allele specificity in counter allele cell lines, c.-26A candidates induced minor counter-allele knockdown (17.5 ± 1.7%, 10 ± 4.9%, 14.5 ± 2.9%, 11.2 ± 3.5%), as did c.-26G candidates (26.5 ± 16%, 14 ± 11.7%, 19.2 ± 14.8%, 30 ± 17.9%, 20.2 ± 20.3%, -5 ± 10.2%, 13.5 ± 18.5%; **Figure 3C–D**), respectively.

**Figure 3.**
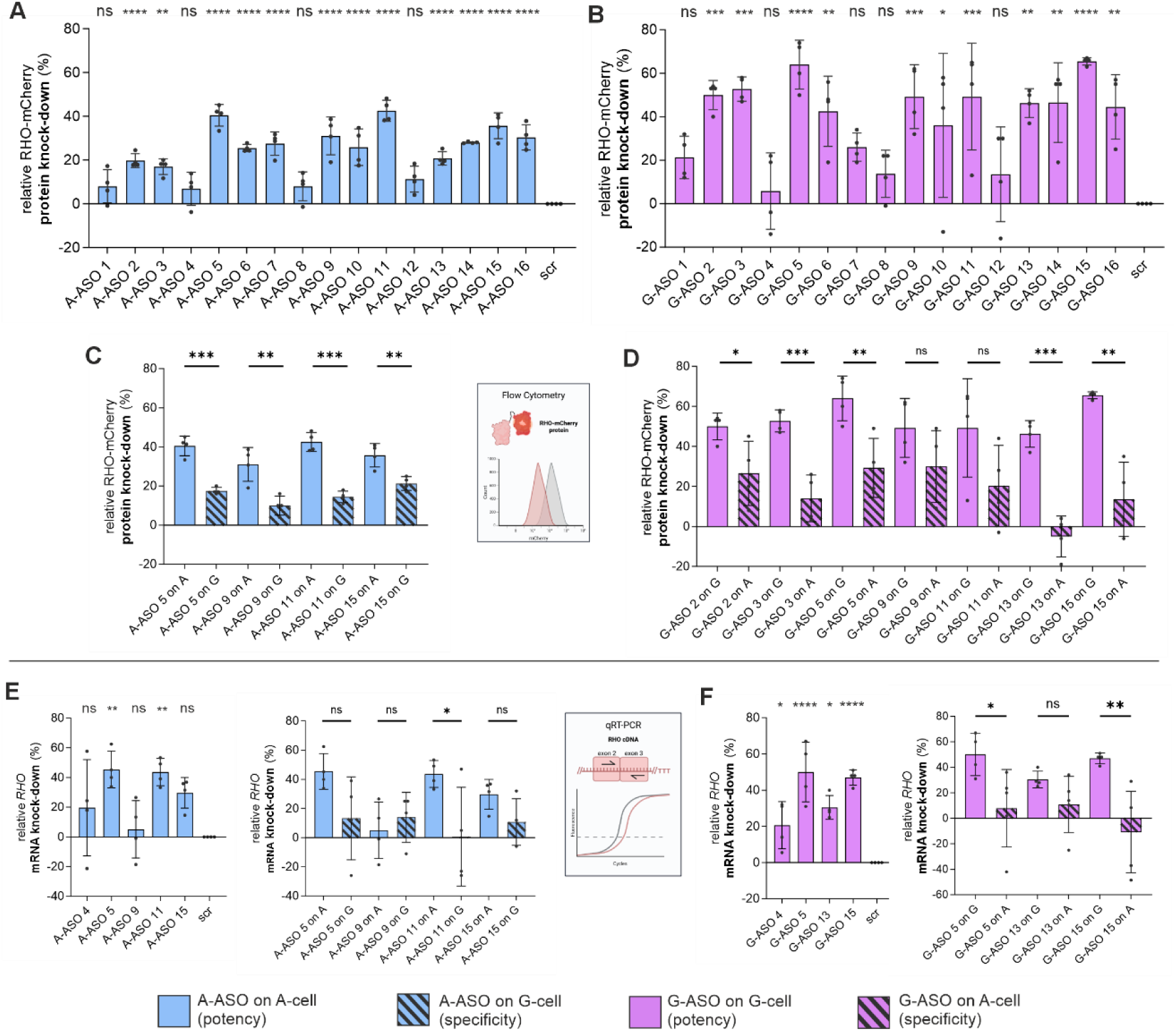
Screening of ASOs in RHO–mCherry reporter cell lines. (A,B) Potency screening for on-target, allele-specific protein knockdown. Sixteen (A) c.-26A-targeting ASOs and sixteen (B) c.-26G-targeting ASOs were transfected into the corresponding c.-26A and c.-26G reporter cell lines. ASO-mediated protein knockdown was quantified by measuring RHO–mCherry fluorescence intensity by flow cytometry relative to scramble ASO (SCR)-treated controls. (C,D) Specificity screening for off-target protein knockdown of selected high-potency ASOs in the reciprocal reporter cell lines for (C) A-targeting ASOs and (D) G- targeting ASOs. (E,F) Confirmation of protein potency and specificity by mRNA transcript knockdown analysis for the selected (E) c.-26A-targeting ASO and (F) c.-26G-targeting ASO. For panels (A,B), statistical significance was determined using one-way ANOVA followed by Dunnett’s multiple-comparison test against the SCR control. For panels (C–F), statistical significance was determined using a non-paired t-test. *p ≤ 0.05, **p ≤ 0.01, ***p ≤ 0.001. Data are presented as mean ± standard deviation. Individual dots represent individual values from the different biological replicates.

Protein knockdown patterns were corroborated at the transcript level via qRT-PCR (**Figure 3E–F**). Low-performing control ASO_4 expectedly showed minimal transcript knockdown (A- ASO_4: 19.6 ± 32.4%; G-ASO_4: 20.6 ± 13%). Based on combined protein and transcript data, c.-26A ASO_5 and ASO_11, and c.-26G ASO_5 and ASO_15 were selected as final lead candidates for downstream validation.

### Testing lead ASOs in retinal organoids

To evaluate endogenous allele-selective knockdown, we utilized patient-derived iPSC- differentiated retinal organoids (ROs) carrying the class III RHO:p.R135W variant in comparison with a CRISPR-engineered wild-type isogenic control (**Supplementary Figures 3–6**). The R135W RHO protein variant was shown to poorly localize in the outer segment membrane (34, 35). First, we quantified relative allelic transcript abundance via targeted high- throughput cDNA sequencing, determining the ratio between c.-26G and c.-26A alleles.

Initial screening was performed by treating ROs with 5 µM ASO for 10 days under an alternating-day washout regimen (**Figure 4A**). Treatment with c.-26A-targeting ASOs (A- ASO_5 and A-ASO_11) increased the proportion of c.-26G transcript, while c.-26G-targeting ASOs (G-ASO_5 and G-ASO_15) enriched the c.-26A transcript, confirming allele-selective suppression. ASO_5 demonstrated the strongest performance in both configurations, achieving non-target transcript increases of 16.0 ± 4.7% (A-target) and 14.7 ± 11.2% (G- target) (**Figure 4B**).

**Figure 4.**
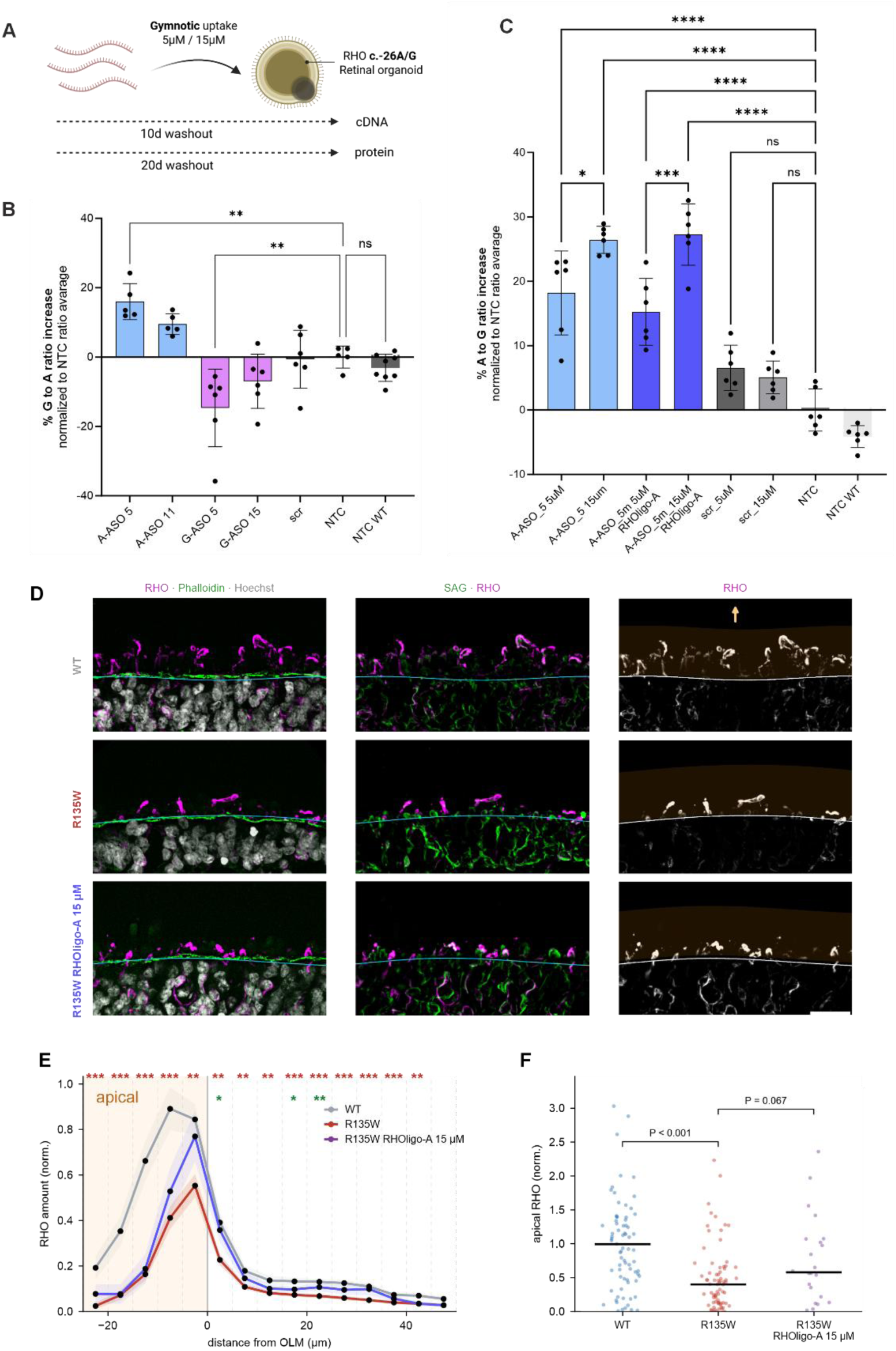
Molecular and phenotypic analysis of lead ASOs in heterozygous iPSC-derived retinal organoids, (A) Experimental outline of the retinal organoid (RO) ASO-treatment. ASOs were added to iPSC-derived retinal organoids for gymnotic uptake at 5 and 15 µM. Following a washout regiment, cDNA analysis was performed after 10 days and protein analysis after 20 days. (B) cDNA illumina sequencing analysis of first batch of ROs differentiated from patient-derived *RHO*:c.403C>T (c.-26A/G *RHO*^R135W/WT^) iPSCs treated with two lead ASOs per SNP (A-ASO_5, A-ASO_11, G-ASO_5, G-ASO_15) and scramble (scr) control. As additional controls served untreated ROs of the same genotype (NTC) as well as ROs derived from the isogenic wildtype control iPSCs (NTC WT). Depicted is the relative ratio of detected c.-26G to c.-26A *RHO* alleles. (C) cDNA illumina sequencing analysis of second batch of ROs differentiated from patient-derived *RHO*:c.403C>T (c.-26A/G *RHO*^R135W/WT^) iPSC treated with A-ASO_5 and methylated A-ASO_5 (A-ASO_5m/RHOlig-A) and scr control at concentrations 5 and 15 µM, respectively. As additional controls untreated ROs of the same genotype (NTC) as well as ROs derived from the isogenic wildtype control iPSCs (NTC WT) were used. (B,C) Statistical significance was determined using ordinary one-way ANOVA *p ≤ 0.05 **p ≤ 0.01, ***p ≤ 0.001, ****p ≤ 0.0001. Data are shown as average ± standard deviation.(D) Representative ROs for WT (c.-26A/G *RHO*^WT/WT^), R135W (c.-26A/G *RHO*^R135W/WT^), and R135W treated with 15 µM RHOligo-A. From left to right: Rhodopsin (RHO, magenta), Phalloidin (green) and Hoechst (grey) merged; SAG (green)-RHO (magenta) merged; and RHO alone (greyscale). Hand-traced outer-limiting membrane (OLM) in cyan. Orange is: apical/segment side; arrow points towards apical/segment direction. Scale bar: 20 µm. (E) RHO amount versus distance from the OLM, in 5 um shells from 25 µm apical (-) to 50 µm inside (+); dotted lines mark shell edges; orange area: apical side. Lines are mean ± SEM. across images, each normalized to its batch-WT peak shell. Red and green asterisks display per-shell significance of the mutant (R135W vs WT) and rescue (R135W A-ASO5 15 uM vs R135W U (F) Apical RHO amount (∼1-26 µm zone apical to the OLM) per image, normalised to the batch-WT median; two-sided Mann-Whitney U. n = 77 WT, 74 R135W and 22 R135W A-ASO5 15 uM images (29, 30 and 11 organoids divided by 2 fully independent experimental batches). Signal was quantified with a fixed per-batch threshold. Statistics: two-sided Mann-Whitney (*P < 0.05, **P < 0.01, ***P < 0.001).

Consequently, we selected A-ASO_5 for further characterization. To potentially reduce immunogenicity (36), we designed and compared in parallel a cytosine-methylated version (A- ASO_5m). Dose-response evaluations at 5 µM and 15 µM revealed no significant performance differences between the two versions of A-ASO_5, indicating no influence of methyl-cytosine modification. At 5 µM, A-ASO_5 and A-ASO_5m achieved non-target transcript increases of 18.2 ± 6.5% and 15.2 ± 5.2%, respectively; at 15 µM, they achieved 26.4 ± 2.1% and 27.3 ± 4.8% (**Figure 4C**). A-ASO_5m was selected as the final lead candidate and renamed RHOligo-A.

Next, RHO protein levels in R135W ROs were compared with those in isogenic WT controls and R135W mutants treated with RHOligo-A at 15 µM (**Figure 4D**; **Supplementary Figure 7**). For WT ROs, first NTC, scrambled ASO treated and RHOligo-A-treated RO were assessed individually (**Supplementary Figure 7A-C**) and then in combination (**Figure 4D-F**); for mutants NTC and scr-treated ROs (**Supplementary Figure 7A-C**) were combined (**Figure 4D-F**). RHO staining level was reduced in R135W mutant organoids not only within the photoreceptor outer segments but also in the inner nuclear layer (INL) (**Figure 4D**). To quantitatively assess the spatial distribution of RHO, the outer edge of the organoid, corresponding to the outer limiting membrane (OLM), was used as a reference point (**Figure 4D**, green). An apical-to-central trajectory was then defined (**Figure 4E**; **Supplementary Figure 7B**), along which consecutive 5-µm shells above and below the OLM were analyzed individually (**Supplementary Figure 7A**). Significant reductions in RHO signal were detected across multiple shells in the mutant organoids (**Figure 4E**, red stars). Notably, RHOligo-A treatment significantly restored RHO levels in three shells (0–5, 15–20, and 20–25 µm), all located within the INL of the organoids (**Figure 4E**). Focusing specifically on the region apical to the OLM, corresponding to the rod photoreceptor outer segments, R135W mutant organoids showed a highly significant reduction in RHO signal compared with WT controls (**Figure 4F**). Treatment with RHOligo-A resulted in a trend toward restoration of RHO levels in this region.

### Safety assessment of RHOligo-A (A-ASO_5m)

To evaluate transcriptome perturbation, wild-type ROs were treated with 15 µM RHOligo-A or a scrambled ASO (SCR) control for 10 days and analyzed via RNA sequencing (RNA-seq). Compared to NTC, we identified 219 and 133 differentially expressed genes (DEGs) for RHOligo-A and SCR, respectively (**Figure 5A**), with 77 overlapping DEGs. On-target activity was confirmed by allele-specific RHO analysis (**Supplementary Figure 8**). Notably, Integrating the 219 RHOligo-A DEGs with *in silico* off-target binding predictions (allowing up to two mismatches/gaps) identified 26 overlapping candidate genes (**Supplementary Table 3**). Multiple identified DEGs (*KRT8, LAMB3, LAMC2, SMN1, RBM47*) are highly enriched in retinal pigment epithelial (RPE) cells, a common RO differentiation by-product. After correcting the transcriptomic analysis for differences in RPE content between individual ROs, all 26 predicted off-target candidates were resolved, indicating that the initial transcriptome changes were driven by differences in RPE abundance rather than sequence-specific off-target effects of RHOligo-A.

**Figure 5.**
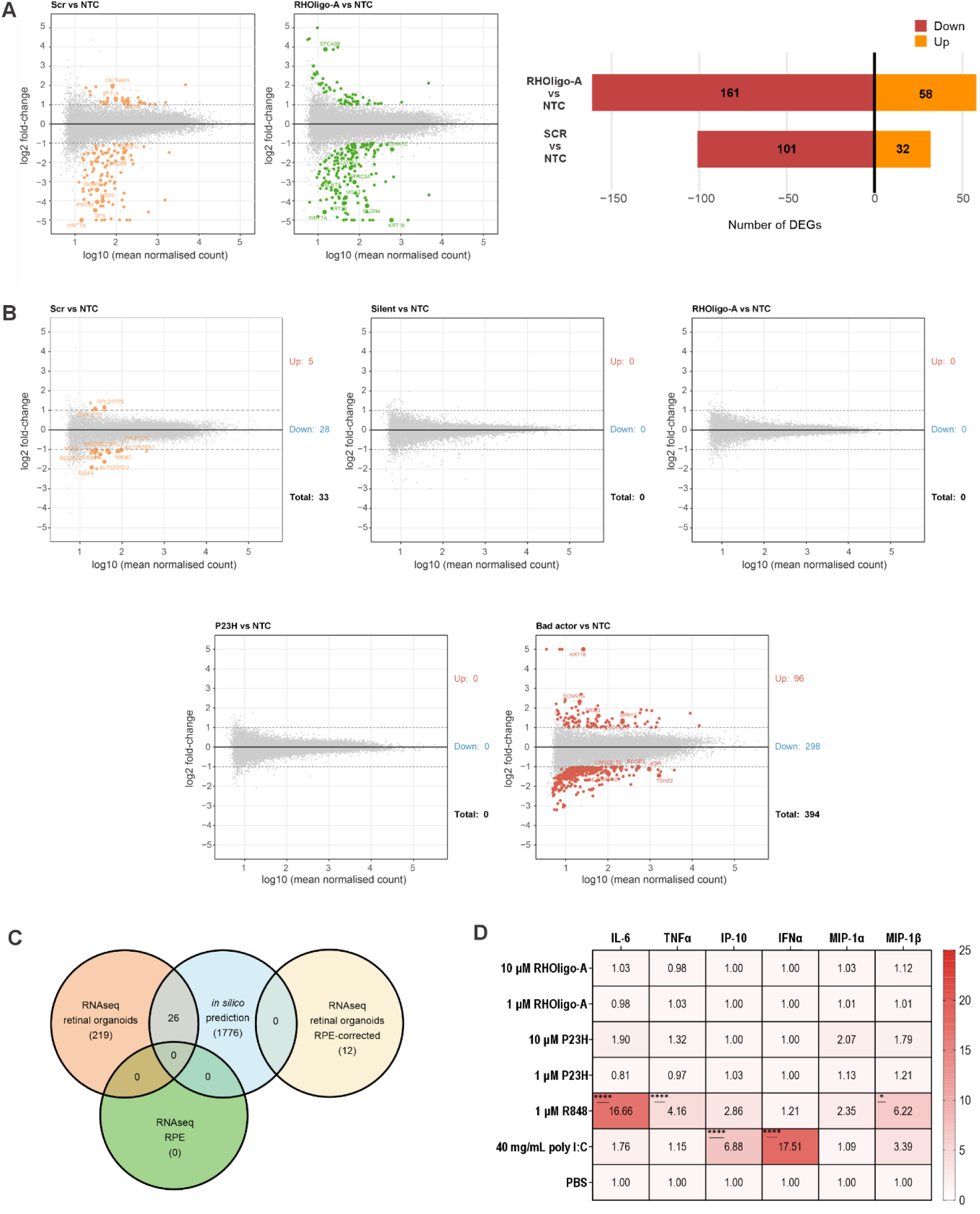
Safety profile and transcriptome assessment of RHOligo-A. (**A**) MA plots displaying differentially expressed genes (DEGs) in retinal organoids (ROs) treated with either a scramble control ASO (SCR) or the lead candidate RHOligo-A, compared against non-treated controls (NTC). The total number of identified DEGs before *in silico* off-target filtering is indicated. (**B**) MA plots of DEGs in iPSC-derived retinal pigment epithelial (RPE) cells following treatment with SCR, a silent RHOligo ASO control (Silent), RHOligo-A, a NHP-validated reference ASO (P23H), and a positive control gapmer known to induce widespread transcriptome disruption (Bad actor). (**C**) Venn diagram intersecting DEGs across all experimental and computational layers, demonstrating 0 overlapping off- target candidates. (**D**) Quantitative cytokine and chemokine release profiles following human PBMC stimulation. The heatmap depicts the geometric mean fold-change relative to PBS control. Statistical significance was assessed via ordinary one-way ANOVA followed by Dunnett’s multiple comparisons post-hoc test, with each treatment condition compared directly back to PBS (*p ≤ 0.05, **p ≤ 0.01, ***p ≤ 0.001 and ****p ≤ 0.0001).

To validate these RPE-associated findings, wild-type iPSC-derived RPE cells (which lack endogenous *RHO* expression) were treated with 15 µM ASO for 3 days and RNA-seq was performed. We benchmarked RHOligo-A against three controls: (i) a known mouse "bad actor" ASO (positive control); (ii) a non-human-primate-validated intravitreal ASO targeted to RHO:p.P23H (P23H) (37); and (iii) a gapless, fully MOE-modified "silent" version of RHOligo- A to isolate chemistry-specific effects (30). As expected, the bad actor induced substantial transcriptional perturbation (394 DEGs). In contrast, neither RHOligo-A, the silent control, nor the P23H control induced any DEGs compared with the NTC, whereas the SCR control resulted in 33 DEGs (**Figure 5B**). Ultimately, intersecting the RO and iPSC-RPE datasets with *in silico* predictions revealed 0 overlapping off-target genes (**Figure 5C**).

To assess RHOligo-A immunostimulatory potential, peripheral blood mononuclear cells (PBMCs) from nine healthy donors were exposed to 1 or 15 µM RHOligo-A. PBMCs from nine healthy donors were exposed to 1 and 10 µM RHOligo-A, and release of IL-6, TNF-α, IP-10, IFN-α, MIP-1α, and MIP-1β was quantified. R848 and poly(I) were used as positive controls, and P23H as a reference control (38, 39). While positive controls induced robust cytokine release, RHOligo-A elicited no significant secretion of any analyte at either concentration. The reference p.P23H ASO showed a modest immune response at 15 µM (**Figure 5D**). Collectively, these findings demonstrate a highly favorable *in vitro* safety profile for RHOligo- A.

### Development of a novel fully humanized *RHO*:c.-26A/G mouse model carrying RHO:p.P347L in *cis* with c.-26A

To evaluate RHOligo-A in a disease-relevant context, humanized mouse lines incorporating the heterozygous *RHO*:c.-26A/G SNP background were generated. We focused on the *RHO*:c.1040C>T(p.P347L) variant, highly prevalent in European patients. This single amino acid substitution is located in the C-terminal cytoplasmic tail of RHO within the VxPx trafficking motif (VAPA in humans), which is essential for trafficking RHO to the rod outer segment. Mislocalization of RHO:p.P347L leads to its accumulation in the inner segment, triggering endoplasmic reticulum (ER) stress and subsequent photoreceptor degeneration (40–43).

Since *RHO*:c.-26A/G resides in the 5′ untranslated region (UTR) of *RHO*, whereas the pathogenic p.P347L variant is located in the 3′ coding region, we pursued full humanization of the mouse *RHO* locus. Two mouse lines were established: a homozygous wild-type line carrying human c.-26G/G hRHO^WT/WT^ and a homozygous mutant line carrying c.-26A/A hRHO^P347L/P347L^. These mouse lines were crossed to generate heterozygous c.-26A/G hRHO^P347L/WT^ offspring for downstream experiments, including natural history characterization (**Figure 6A–B**). *RHO* transcript quantification revealed comparable expression levels between hRHO^WT/WT^ and hRHO^P347L/WT^ lines, though both were reduced relative to C57BL/6J controls (**Figure 6C**). Retinal structural analysis of outer nuclear layer (ONL) thickness showed similar profiles between c.-26G/G hRHO^WT/WT^ and C57BL/6J mice. Overall, retinal structure in c.- 26G/G hRHO^WT/WT^ mice remained stable over the 6-month observation period. In contrast, heterozygous c.-26A/G hRHO^P347L/WT^ animals exhibited marked retinal degeneration as early as 1 month of age, with progressive deterioration over time (**Figure 6D–I**). By 6 months, only a residual ONL was detectable, predominantly proximal to the optic nerve (**Figure 6I**). Electroretinography (ERG) analysis revealed no significant differences between c.-26G/G hRHO^WT/WT^ and C57BL/6J mice up to 6 months of age (**Figure 6J–M**). Conversely, c.-26A/G hRHO^P347L/WT^ mice exhibited a progressive decline in retinal function over time, with reductions in both scotopic maximal responses and photopic cone responses (**Figure 6L–M**). Notably, scotopic rod responses in mutant animals remained consistently reduced but relatively stable throughout the time course, at approximately threefold lower level than in c.-26G/G hRHO^WT/WT^ controls (**Figure 6J**).

**Figure 6.**
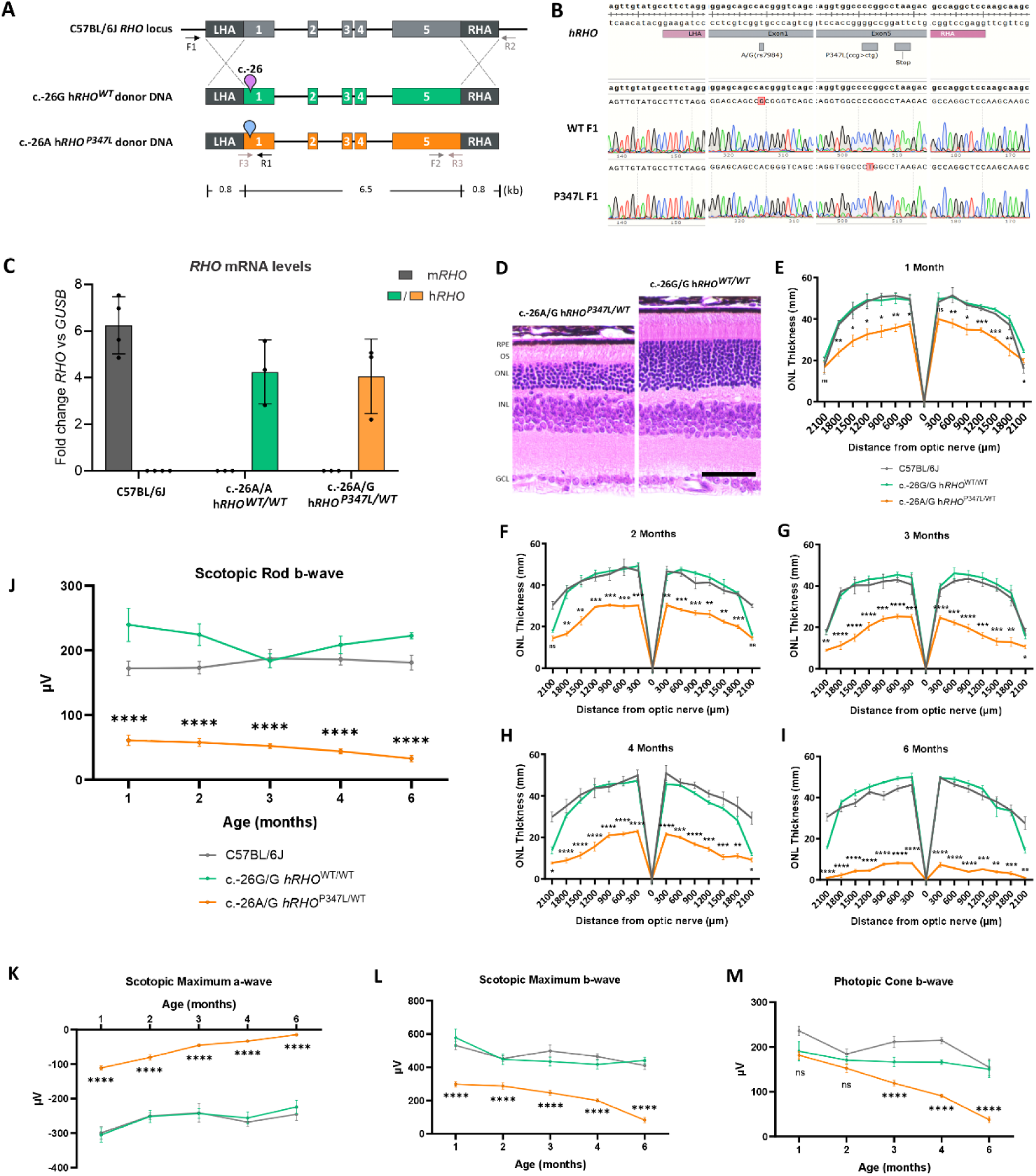
Generation and natural history of humanized c.-26A/G h*RHO*^P347L/WT^ mouse model. (A) Schematic representation of homology-directed-repair (HDR)-mediated insertion of human c.-26G RHO^WT^ and human c.-26A RHO^P347L^ donor DNA into the endogenous *Rho* locus of C57BL6J mice (LHA=Left Homology Arm, RHA = Right Homology Arm). (B) Comparative electropherograms of the humanized *RHO* locus of F1 generation of generated humanized WT and humanized RHO^P347L^ mice. To selectively amplify the human *RHO* locus in F1 generations, human RHO-specific primers were used (C) qRT-PCR with assays specific for the murine and human *RHO* gene with retinal cDNA from animals homozygous for the humanized WT *RHO* allele and heterozygous for the humanized WT and humanized P347L *RHO* allele demonstrating exclusive expression of human *RHO* compared to non-humanized parental mouse line. (D) Representative hematoxylin and eosin–stained retinal sections highlighting the peripheral retinal region located approximately −1800 µm from the optic nerve, comparing humanized heterozygous mutant (hRHO^WT/P347L^, left) and humanized WT (hRHO^WT/WT^, right) at 28 days of age. (E, F, G, H, I) Spider plots showing outer nuclear layer (ONL) thickness of the humanized and parental mouse lines at 1,2,3,4 and 6 months of age. (n = 3-4). Statistical significance was determined using unpaired t-test between humanized c.-26A/G hRHO^P347L/WT^ and c.-26G/G hRHO^WT/WT^. ****p ≤ 0.0001. Data are shown as average ± standard deviation. *p ≤ 0.05, **p ≤ 0.01, ***p ≤ 0.001, ****p ≤ 0.0001, ns = not significant. (J, K, L, M) Scotopic rod b-wave, scotopic maximum a-wave, scotopic maximum a-wave and photopic cone b-wave amplitudes of the humanized and parental mouse lines, measured at 1,2,3,4 and 6 months of age, respectively (n = 3-28). Statistical significance was determined using ordinary one-way ANOVA. ****p ≤ 0.0001. Data are shown as average ± standard deviation. (C) Individual dots represent individual values from the different biological replicates.

### Testing lead A-ASO_5m (RHOligo-A) in humanized RHO mouse model

First ocular tolerability of RHOligo-A was assessed by unilateral intravitreal injections of low (LD, 10 µM) or high (HD, 50 µM) dose in 4-week-old C57BL/6 mice. Retinal function evaluated 30 days post-injection (PI) via ERG showed no significant alterations (**Supplementary Figure 9A–B**), indicating good ocular tolerability and supporting evaluation in the newly established c.-26A/G hRHO^P347L/WT^ mouse model.

RHOligo-A was subsequently assessed for *in vivo* target engagement and functional efficacy in the mouse model. c.-26A/G *RHO*^P347L/WT^ mice aged P21–P27 received single unilateral intravitreal injections in the right eye (LD or HD), while contralateral left eyes were mock-treated with PBS (**Figure 7A**). Ten days PI, bulk *RHO* transcript reduction reached 30 ± 5% (LD) and 41 ± 11% (HD), whereas c.-26G/G hRHO^WT/WT^ mice exhibited minimal knockdown (6 ± 18% LD; 11 ± 10% HD) (**Figure 7B**). Allele-preferential targeting in *vivo* was confirmed by determining the relative abundance of c.-26A– and c.-26G–containing transcripts via next generation sequencing (NGS) and integrating these data with bulk knockdown measurements to determine allele-specific suppression (**Supplementary Table 4**). This analysis demonstrated a strong in vivo preference of RHOligo-A for the c.-26A allele, with knockdown levels of 51 ± 9% (for the c.-26A allele) vs. 2 ± 16% (for c.-26G) at LD and 64 ± 3% (for c.- 26A) vs. 15 ± 4% (for c.-26G) at HD (**Figure 7C**).

**Figure 7.**
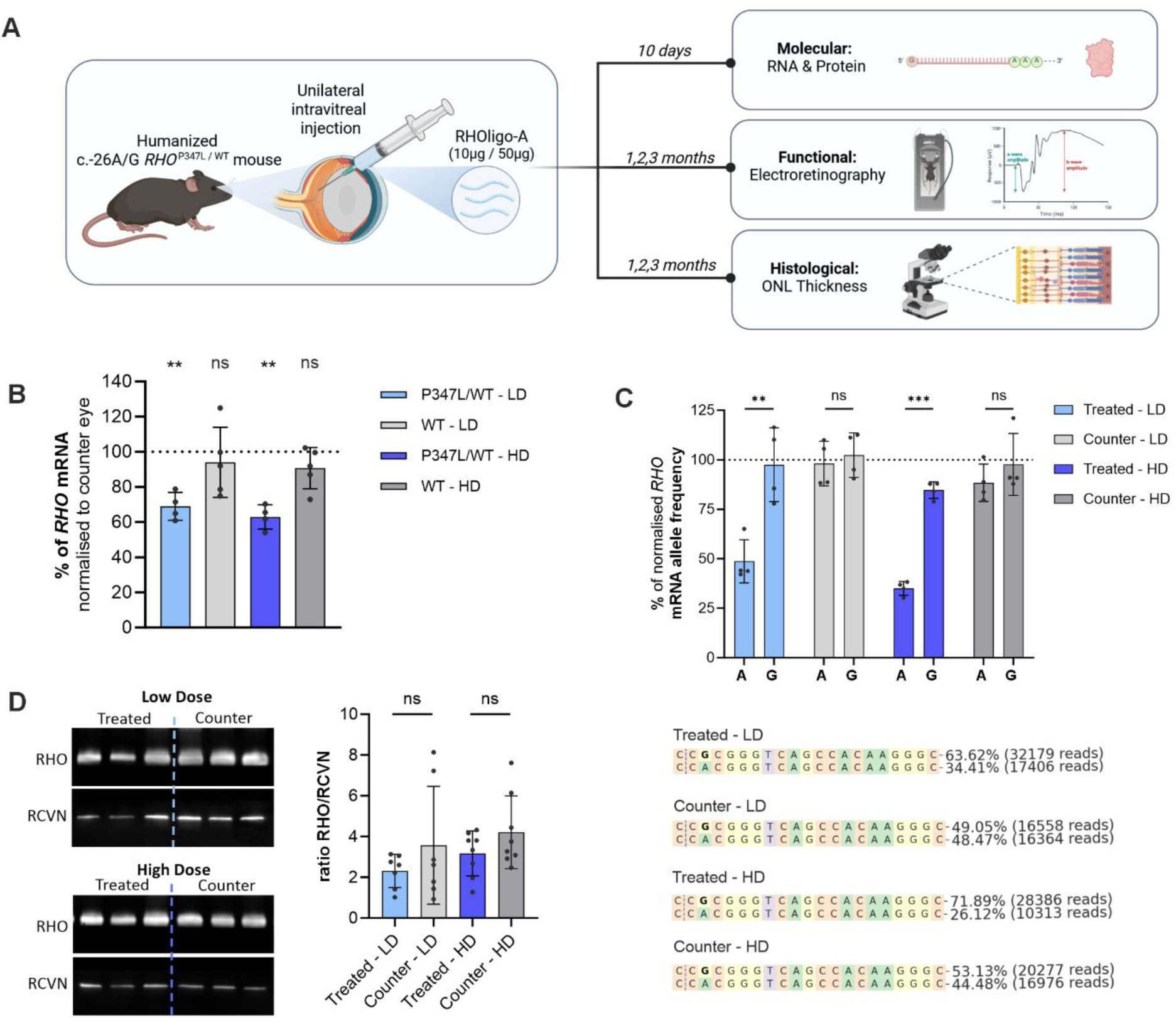
Experimental outline and molecular analysis of RHOligo-A treatment in the humanized c.-26A/G RHO^P347L/WT^ mouse model. (A) Outline of the in vivo experiments performed, timelines, and read outs analyzed (B) qRT–PCR analysis of total (bulk) *RHO* transcripts. Data from treated eyes were normalized to the contralateral PBS-treated eyes. *Pde6g*, a rod-specific gene, was used as the reference control. (C) Allele-specific *RHO* knockdown was quantified by determining the relative abundance of c.-26A– and c.-26G–containing *RHO* transcripts using next-generation sequencing (NGS) normalized to the bulk *RHO* knockdown levels. Examples for NGS results from one low dose- and one high dose-treated animal is depicted (**Supplementary Table 4**) (D) Immuno blot analysis of total RHO protein levels in RHOligo-A-treated and contralateral eyes following low-dose (LD) and high-dose (HD) injection. Recoverin (RCVN) was used as a loading control. Semi-quantitative band densitometry plot analysis is shown for n = 7 mice (LD) and n = 8 mice (HD). (B,C,D) Statistical significance was determined using a paired t-test. **p ≤ 0.01, ***p ≤ 0.001. Data are shown as average ± standard deviation. Individual dots represent individual values from the different biological replicates.

Semi-quantitative immunoblot analysis revealed a subtle trend toward reduced total RHO protein in treated eyes compared to controls at both doses (**Figure 7D**).

Retinal function was assessed by ERG at 1-, 2-, and 3-months PI. Unilateral intravitreal PBS injections served as procedural controls; no differences between injected and contralateral eyes were observed at 1 month PI, indicating no effect on retinal function (**Supplementary Figure 10A**). At 1 month PI, the HD group showed robust scotopic rod b-wave restoration in treated eyes compared to controls (147 ± 15 µV vs. 65 ± 14 µV), reaching near-WT levels and even exceeding the baseline response at P30 (**Figure 8A–B**). For the LD group, scotopic rod-specific b-wave responses were 117 ± 15 µV in treated eyes vs. 48 ± 6 µV in controls (**Figure 8B**). At 2 months PI, scotopic b-wave responses for both treatment groups normalized to approximately 100 µV, representing average increases over PBS-treated eyes of 226 ± 25% (LD) 211 ± 20% (HD). At 3 months PI, scotopic b-wave responses remained elevated, with average percentage increases over controls of 174 ± 24% (LD) and 189 ± 23% (HD) (**Figure 8C**). Significant enhancement in the scotopic maximum response was also significant across all three time points for both doses (**Figure 8D–E**). Expectedly, no differences in photopic cone response were detected between treated and control eyes within this time frame (**Figure 8F**). In c.-26G/G hRHO^WT/WT^ mice, the HD caused a modest reduction in scotopic rod b-wave and maximal responses, whereas the LD group showed no significant differences (**Supplementary Figure 10B–C**).

**Figure 8.**
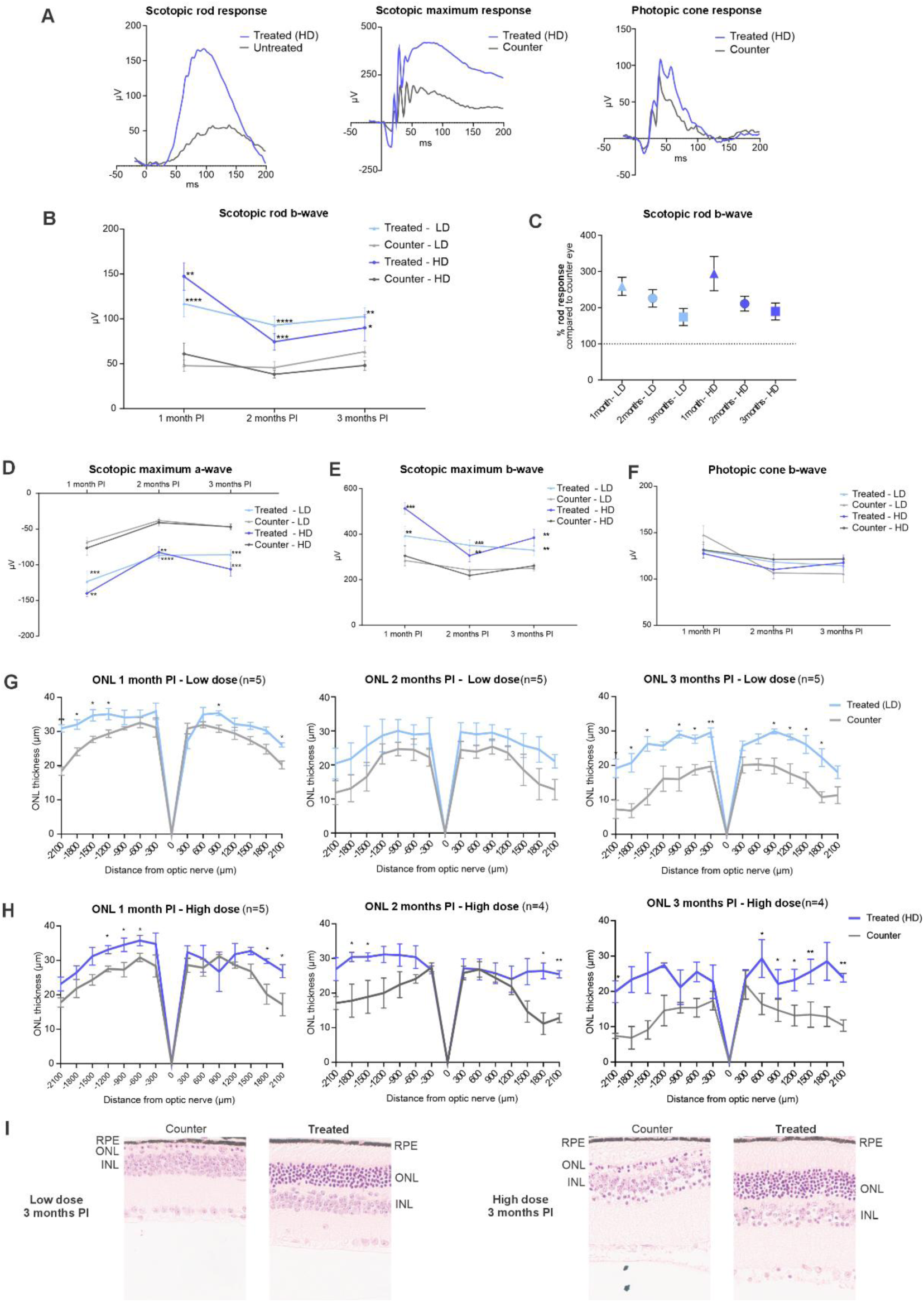
Retinal function and structural preservation following RHOligo-A treatment in the humanized *c.-26A/G RHO^P347L/WT^* mouse model. (A) Representative ERG traces recorded 1-month post-injection (PI) from a mouse treated with the high dose (HD), showing scotopic rod, scotopic maximum, and photopic cone responses. (B) Scotopic rod b-wave amplitudes measured over the three-month period for low-dose (LD) and high-dose (HD) treated groups. (C) Percentage increase in scotopic rod b-wave responses in treated eyes compared with contralateral PBS-treated control eyes at each of the three time points for both LD and HD. (D) Scotopic maximum a-wave amplitudes, (E) scotopic maximum b-wave amplitudes, and (F) photopic cone b-wave amplitudes recorded over the three-month period for LD and HD groups. (G, H) Spider plots showing outer nuclear layer (ONL) thickness for the (G) LD and (H) HD groups over the three-month treatment period. (I) Representative hematoxylin and eosin–stained retinal sections highlighting the peripheral region located approximately −1800 µm from the optic nerve, comparing LD-treated (left) and HD-treated (right) eyes with their corresponding contralateral PBS-treated controls. Statistical significance was determined using a paired t-test. *p ≤ 0.05, **p ≤ 0.01, ***p ≤ 0.001. Data are shown as average ± standard error of mean. For (B,C,D,E,F) n = 5-14 per group (Supplementary Table 5).

Preferential knockdown of mutant *RHO* transcript resulted in substantial preservation of retinal architecture, as demonstrated by sustained ONL thickness throughout the 3-month observation period (**Figure 8G–I**). Protective effects were detectable at 1 month PI for both doses and increased progressively over time, reaching maximal separation from controls at 3 months. Structural preservation was most pronounced in the peripheral retina; at 3 months PI, treated eyes retained approximately 6–8 rows of photoreceptor nuclei, whereas control eyes were frequently reduced to a residual nuclear layer with single nuclei (**Figure 8I**).

To assess the effect of RHOligo-A on the rate of degeneration, quantitative aggregate analysis of ONL thickness was performed using the area under the curve of the spider plot. Results indicate that 79.15 ± 2.99% and 79.01 ± 5.79% of retinal structure was preserved between months 1 and 3 PI in treated eyes at LD and HD, respectively, compared to 54.53 ± 3.89% and 52.77 ± 5.70% in controls (**Supplementary Figure 11**). Consistently, the relative decline in aggregate ONL thickness between months 2 and 3 was limited to 3.96 ± 6.84% (LD) and 11.78 ± 6.73% (HD) in treated eyes versus 24.28 ± 8.6% and 33.42 ± 8.05% in controls.

Ultrastructural analysis by electron microscopy of HD-treated c.-26A/G hRHO^P347L/WT^ mice at 3 months PI demonstrated preserved retinal architecture, featuring clearly identifiable, structurally intact photoreceptors and well-organized, functional ribbon synapses. In contrast, PBS-treated eyes showed complete degeneration of the photoreceptor layer, leaving no photoreceptor cells or organized synaptic structures (**Supplementary Figures 12–13**).

## DISCUSSION

Genotyping across multiple RHO-adRP cohorts showed that the benign *RHO*:c.-26A/G SNP is frequently heterozygous, providing a genetic entry point for allele discrimination and enabling mutation-independent, allele-specific therapeutic strategies. This paradigm addresses a major limitation of mutation-specific approaches for RHO-associated disease, where extensive allelic heterogeneity constrains patient reach and translational scalability (6, 11). Guided by this rationale, we developed ASOs capable of single-nucleotide discrimination between the two SNP alleles. Because gapmer ASOs typically induce robust but incomplete transcript suppression, limited activity on the non-target allele is unlikely to have clinically relevant consequences. In contrast, maintaining suppression of the dominant toxic allele is expected to provide a substantially greater therapeutic benefit than the potential risk associated with partial reduction of total rhodopsin levels (13, 14, 44). Indeed, heterozygous carriers of null alleles are asymptomatic, whereas homozygous carriers and *Rho*^-/-^ mice exhibit a severe phenotype, indicating a functional window between haplosufficient and null states (13, 14, 45). However, the lower boundary of safe RHO abundance remains undefined.

To prioritize candidate ASOs, we established RHO–mCherry reporter cell lines as a functional screening platform. Protein-level readouts of potency and specificity by flow cytometry demonstrated high reproducibility and low inter-replicate variability compared to transcript-based measurements, providing a robust proxy for knockdown activity. Owing to differences in basal *RHO* expression between the c.-26A and c.-26G reporters, this platform is best suited for comparative ranking of molecules within each line rather than quantitative comparisons between lines. Interestingly, the panel of A- and G-ASOs yielded similar relative potencies, suggesting that a single nucleotide change (either A or G) does not substantially affect molecular target engagement of a fully complementary ASO at this site.

Endogenous knockdown was validated in iPSC-derived retinal organoids. Due to substantial variability in RHO expression across organoids, allele-selective activity was quantified via relative enrichment of the non-target allele using targeted cDNA sequencing rather than conventional quantitative PCR normalized to housekeeping genes. Through this approach, A- ASO_5 (RHOligo-A) and G-ASO_5 emerged as lead candidates, consistently inducing preferential suppression of the targeted allele. Notably, a 5-methyl-cytosine-modified version of RHOligo-A retained equivalent potency and dose responsiveness, supporting the compatibility of chemical optimization strategies to improve translational properties without compromising efficacy.

The RHO:p.R135W variant has been reported to exhibit impaired plasma membrane localization; however, whether the mutant protein also affects WT RHO localization in the heterozygous context remains unclear (34, 35). Following validation of endogenous allele-preferential knock-down of RHOligo-A in retinal organoids, we used the polarized distribution of RHO within photoreceptors, delineated by phalloidin staining (OLM), as a phenotypic readout. RHO localization differed significantly between R135W mutant and isogenic WT organoids, suggesting that the mutant protein may at least partially impair WT RHO localization or expression. Notably, a single 15 µM dose of RHOligo-A led to a clear trend towards restoration of RHO localization towards the WT pattern after 20 days. The rapid emergence of a WT-like RHO localization pattern following RHOligo-A treatment, despite the short exposure period and advanced maturation of the ROs, supports its potential to correct disease-relevant phenotypes associated with Class III RHO mutations.

Safety assessments based on off-target transcriptome evaluations in ROs and iPSC-derived RPE cells, integrated with *in silico* prediction, alongside cytokine release assays in PBMCs, indicated a favorable safety profile for RHOligo-A. Regarding transcriptome perturbation, no DEGs were nominated when integrating multilayer analysis of ROs, iPSC-RPE cells and *in silico* predicted off-targets, suggesting a low transcriptome perturbation potential of RHOligo-A. Notably, different overlapping DEGs between RHOligo-A and SCR were identified in ROs, suggesting sequence-independent effects such as protein interactions mediated by ASO chemistry, which will have to be assessed *in vivo* through dedicated toxicology studies (46, 47). In PBMC, while the lead RHO:p.23H ASO validated in non-human primates induced a modest increase in cytokine release (IL-6, MIP-1α, and MIP-1β) at 10 µM, RHOligo-A did not elicit detectable cytokine responses under the conditions tested, further derisking its immunogenetic potential (48, 49).

To evaluate the translational potential *in vivo*, we generated a genomically fully humanized c.- 26A/G hRHO^P347L/WT^ mouse model, which recapitulates both disease biology and the clinically relevant SNP configuration. In this model, RHOligo-A achieved robust, allele-preferential transcript suppression along with sustained functional and structural retinal preservation. Interestingly, transcript knockdown did not proportionally reduce total RHO protein levels. Plausibly, selective suppression of mutant RHO promotes stabilization and improved trafficking of the wild-type protein, increasing its steady-state abundance and offsetting the expected reduction in total RHO (50, 51). Furthermore, the reduced ERG responses in high- dose hRHO^WT/WT^ mice likely reflect off-target *RHO* knockdown rather than toxicity, consistent with the unaltered ERG profiles observed during C57BL/6J tolerability testing.

Although intravitreal clearance is faster in mice than in humans, and rodent doses may therefore overestimate the concentration required clinically, formal ocular pharmacokinetic and toxicology studies will be needed to define the human starting dose and dosing interval (52). The translational framework for this modality is further reinforced by clinical experience with QR-1123, a gapmer ASO targeting RHO:p.P23H developed by Ionis Pharmaceuticals, which demonstrated favorable ocular safety and early efficacy signals in phase 1/2 clinical studies (NCT04123626) (37).

Collectively, our findings establish the SNARE platform as a broadly applicable, mutation-independent therapeutic strategy for RHO-adRP. Evidence of phenotypic rescue potential was provided across two distinct *RHO* mutations belonging to different mechanistic classes, supporting the broad applicability of this approach. Validation across reporter systems, patient-derived retinal organoids, and a humanized *in vivo* mouse model consistently demonstrated allele-selective suppression alongside molecular, structural, and functional rescue *in vivo*. Based on our cohort analyses, RHOligo-A alone can address approximately 15.4% of patients with RHO-associated disease, while further exploring the complementary G-ASO_5 strategy could expand coverage to roughly 28.5%. Given population-specific rs7984 allele frequencies in gnomAD v4. 1. 1, the clinical reach may be even greater in certain ethnic groups (e.g., a heterozygous allele frequency of 36% in East Asian populations). Overall, these findings provide strong preclinical proof of concept for allele-selective, mutation-independent ASO therapies in RHO-adRP and other gain-of-function IRD, establishing RHOligo-A as our lead candidate for future clinical development.

## METHODS

### Plasmid cloning

The parental transfer *RHO*-mCherry lentivirus plasmids (REV_PGK-RHO-Linker-mCherry- P2A-ZeoR-WPRE) was cloned by digesting the pKLV2.2-mU6gRNA5(SapI)- hU6gRNA5(BbsI)-PGKpuroBFP-W (Addgene, catalog no. 72667) lentiviral transfer plasmid by *Apa*I and *KpnI* and cloning the PGK-RHO-mCherry-P2A-ZeoR-WPRE fragments by NEBuilder® HiFi DNA Assembly Master Mix (NEB). Specifics primers used are listed in **Supplementary Table 2**. The c.-26G and c.696+4C variants were introduced in the parental plasmid by in vitro mutagenesis using PfuUltra II Fusion High-fidelity DNA Polymerase (Agilent), resulting in the transfer plasmids containing c.-26A+696+4T and c.-26G+696+4C SNPs. Plasmid sequences are provided in **Supplementary Table 6**.

### Gapmer antisense oligonucleotide

For *in vitro* screening, c.-26A-targeting ASOs were synthetized by ProQR (Leiden, NL) while c.-26G-targeting ASOs were purchased from Eurogentec. Low endotoxin level ASO (<0.5 EU/mg) for retinal organoids, PBMC assay and, *in vivo* experiments were purchased from LGC Axolabs GmbH.

### Patient genotyping and mutation phasing

Genomic DNA samples from *RHO*-adRP patients were first genotyped for the presence of the target heterozygous SNP. Approximately 50–100 ng of genomic DNA was amplified using Taq DNA Polymerase S (Genaxxon Bioscience), and PCR products were analyzed by Sanger sequencing. Samples identified as heterozygous for rs7984 were subsequently phased to determine which SNP allele was in cis with the pathogenic *RHO* variant. Phasing was performed using a two-step allele-specific PCR strategy. Briefly, the full *RHO* locus was first amplified using KAPA HiFi HotStart ReadyMix (Roche) with a forward primer incorporating LNA chemistry overlapping the c.-26G allele and a common reverse primer, enabling preferential amplification of the c.-26G allele over c.-26A. In a second PCR reaction, exon-specific primers were used, depending on the location of the patient’s *RHO* variant, with the first-round product serving as template. The resulting amplicons were Sanger sequenced. Detection of the pathogenic variant in this allele-selective amplicon indicated that the mutation was in cis with the c.-26G allele, whereas detection of the wild-type sequence indicated cis association of the *RHO* variant with the c.-26A allele. List of primers provided in **Supplementary Table 2**.

### Sanger sequencing

PCR and RT-PCR amplicons yielding a single product were treated with ExoSAP-IT^TM^ (Applied Biosystems) according to the manufacturer’s instructions to remove residual primers and nucleotides. For amplification reactions producing multiple bands, individual fragments were excised from agarose gels and purified using standard gel extraction procedures. ExoSAP-treated products or 20 ng of gel-purified DNA were sequenced using the SupreDye v3.1 Cycle Sequencing Kit (EdgeBio) following the manufacturer’s protocol. Sequencing reactions were analyzed on an ABI PRISM 3130xl Genetic Analyzer (Thermo Fisher Scientific). Plasmid constructs were fully sequenced using Oxford Nanopore long-read sequencing (Microsynth AG). Resulting sequence data were aligned and analyzed using the Benchling platform.

### Targeted high-throughput sequencing

Illumina adapter sequences were introduced by PCR amplification of target amplicons using degenerate adapter-containing primers. A second PCR was performed to incorporate unique sample barcodes. Barcoded amplicons were pooled at approximately equimolar ratios based on densitometric quantification from agarose gel images. The pooled library was purified using AMPure XP beads (Beckman Coulter) and quantified by Qubit fluorometric measurement (Thermo Fisher Scientific). Libraries were sequenced by Novogene using an Illumina platform. Allele-frequency analysis of sequencing data was performed using CRISPResso2 (53).

### Cell lines and culture conditions

HEK293T and reporter cells were cultivated in Dulbecco’s Modified Eagle Medium (DMEM) supplemented with 10% Fetal bovine serum (FBS, Thermo Fisher Scientific), 10 U/mL penicillin–streptomycin (Thermo Fisher Scientific) and 2.5 µg/ml amphotericin B (Capricorn Scientific). RHO*-*mCherry reporter cell lines were moreover cultivated with 100 µg/ml zeocin (InvivoGen) at 37°C in a 5% CO_2_ humidified atmosphere.

A skin biopsy was obtained from a c.403C>T p.R135W *RHO*-adRP patient after written informed consent, in accordance with institutional guidelines and with approval from the local ethics committee (project no. 124/2015BO1). Primary dermal fibroblasts were isolated from the biopsy, expanded, and maintained in DMEM supplemented with 20% FBS and 10 U/mL penicillin–streptomycin (hereafter referred to as fibroblast medium). Cells were cultured under standard conditions at 37 °C in a humidified atmosphere containing 5% CO₂.

iPSCs were cultivated in Essential 8 (E8) Flex medium (Gibco) with puromicin (InvivoGen) and routinely passaged using ReLeSR (STEMCELL Technologies) at a 1:6 split ratio plated onto Matrigel-coated 10-cm dishes (STEMCELL Technologies) and incubated at 37 °C in a humidified 5% CO₂ incubator.

Retinal pigment epithelial (RPE) cells were differentiated from iPSCs derived from a genetically screened human donor (National Eye Institute, USA; Prof. Bharti laboratory) (54) and characterized previously (cell line 2) (55). iPSCs were maintained on hESC-Matrigel (Corning) coated plates in E8 medium with daily changes. Differentiation was performed according to a described protocol (56). Briefly, at 70–80% confluency, iPSCs were switched to differentiation medium (Gibco: DMEM/F12, NEAA, KnockOut SR) replaced every 2 days for 42 days with specific stage-formulations: 10 mM nicotinamide (Days 0–7, Sigma-Aldrich), 100 ng/ml activin A (days 7–14; Peprotech), and 3 µM CHIR99021 (days 14–42, Tocris). At day 42, cells were enriched in medium containing ROCK inhibitor (StemMACS Y27632) and seeded onto Geltrex (Gibco) or hESC-Matrigel coated plates for expansion. Cells were matured for at least 30 days with twice-weekly medium changes, and passaged using TrypLE (Thermo Fisher Scientific). Experiments were performed using passage 2–3 cells cultured in 48-well plates for at least 50 days prior to treatment.

### Establishment of the RHO-mCherry reporter cell lines

To produce the *RHO-mCherry* lentivirus particles, the cloned transfer plasmids, the packaging psPAX2 plasmid (Addgene #12260) and the envelope pMD2.G plasmid (Addgene #12259) were co-transfected using Lipofectamine® 3000 into HEK293T cells at a molar ratio of 4:3:1, respectively, according to the manufactureŕs protocol. In brief, 8x106 cells were seeded in 12 ml of DMEM +10% FBS, plated in a 10-cm dish, and incubated overnight at 37 °C in a 5% CO2 humidified atmosphere. The following day, the cells were transfected with a total amount of 15 μg of plasmids in a total volume of 6 ml fresh DMEM +10% FBS without antibiotics. 7 h post-transfection the medium was replaced with 12 ml DMEM containing 2% FBS. 72 h post-transfection, the supernatant was harvested, filtered through a 0,45 μm filter and concentrated via ultracentrifugation in a 20% sucrose cushion at 27,000xg for 2 h at 4°C. The pellet containing the lentiviral particles was carefully resuspended in 150 μl Opti-MEM and stored at -80 °C.

To generate stable transgenic cell lines, 1x105 HEK293T cells were seeded in a well of a 24- well plate in 500 µl polybrene culture medium (DMEM + 10%FBS + 4 µg/ml Polybrene (SantaCruz)). 50 µl of the lentivirus preparation was added, mixed and the plate incubated at 37°C and 5% CO2. Twenty-four hours later, the medium was replaced by fresh DMEM + 10% FBS and 72 h post-transduction, the cells were put under selective pressure by addition of the selection antibiotic zeocin at a final concentration of 100 µg/ml. After expansion to a 6-well plate, cells were singularized using Accutase (Gibco) and low-density plated into 6 cm-dishes. A few days later, grown colonies derived from single clones were picked and transferred into a 24-well plate with medium containing zeocin and expanded for the following days. Clones for further characterization were selected based on high mCherry expression as determined by fluorescence microscopy.

### Generation of induced pluripotent stem cell lines

Patient-derived *RHO*:c.403C>T p.R135W (c.-26A/G RHO^R135W/WT^) iPSCs were generated from fibroblasts by reprogramming with the Yamanaka factors (57, 58). Briefly, 3 µg of plasmids encoding OCT3/4, SOX2, KLF4, L-MYC, and LIN28 (in equimolar amounts) were transfected into 5 × 10⁵ fibroblasts using the NEON electroporation system (Thermo Fisher Scientific) with 100 µL electroporation tips. Electroporation parameters were set to 1,650 V, 10 ms pulse width, and three pulses. After transfection, cells were cultured for 2 days in fibroblast medium supplemented with 2 µg/L fibroblast growth factor-2 (FGF-2; PeproTech). The medium was then replaced with E8 Flex medium (Thermo Fisher Scientific) containing 100 µM sodium butyrate (Sigma-Aldrich). Seven days post-transfection, cells were dissociated with Trypsin-EDTA (Thermo Fisher Scientific) and 1 × 10⁵ cells were plated onto Matrigel-coated 10-cm dishes (STEMCELL Technologies). Emerging iPSC colonies were manually picked and expanded on Matrigel-coated six-well plates in E8 Flex medium.

An isogenic WT iPSC line for the patient-derived *RHO*:c.403C>T iPSC line was generated by CRISPR/Cas9-mediated knock-in. A single guide RNA (sgRNA - IDT) and an asymmetrical single-stranded oligodeoxynucleotide (ssODN - IDT) donor template was designed to correct the *RHO* mutation on the Benchling platform (**Supplementary Table 2**). Briefly, iPSCs were washed with PBS and dissociated using accutase (Gibco). After centrifugation, cells were resuspended in R buffer. The sgRNA/*Sp*Cas9 RNP complexes (250 pmol:sgRNA and 125 pmol SpCas9) (IDT) were assembled and mixed with 500,000 cells along with 400 pmol ssODN (IDT) and used for electroporation with a Neon electroporation system. Electroporation was performed using the NEON transfection system according to the manufacturer’s instructions at 1400V 4ms and 2 pulses. Following electroporation, cells were plated onto vitronectin-coated (Thermo Fisher Scientific) dishes and cultured under standard iPSC conditions until confluent. Then, iPSCs were dissociated into single cells, and approximately 1 × 10⁴ cells were seeded per vitronectin-coated 10-cm dish. Emerging colonies were manually picked, transferred to 24-well plates, and expanded for genotyping and clone selection. Genomic DNA was isolated using the QuickExtract DNA Extraction Solution (Lucigen, Mandel). Correctly edited clones were confirmed by PCR amplification of the target by Taq DNA Polymerase S (Biozym) followed by Sanger sequencing. Primers are listed in **Supplementary Table 2**.

To assess genomic integrity of the selected clone, genomic DNA was extracted using the NucleoSpin Tissue Mini Kit (Macherey-Nagel) according to the manufacturer’s instructions. Genome-wide copy number variation (CNV) analysis was performed by Life & Brain Genomics (Bonn, Germany) using the Infinium OmniExpressExome-8 v1.4 BeadChip microarray (Illumina). Data processing and CNV detection were carried out in GenomeStudio software (Illumina) using the cnvPartition plugin. Genomic integrity and potential chromosomal aberrations were further evaluated through manual inspection of B-allele frequency and log R ratio plots.

### Generation of fully humanized c.-26A/A *RHO^P347L/P347L^* and c.-26G/G *RHO^WT/WT^* mouse models

To create the humanized *RHO* mouse models, the mouse *RHO* locus was cut by dual SpCas9/gRNAs and the gene was replaced with a donor template containing a full-length human *RHO* gene fragment via homology-directed repair HDR. In brief, the human *RHO* DNA from exon 1 to exon 5 and the ∼800-bp upstream and downstream homologous arms fragments were amplified, purified, and then cloned into the pZac2.1 backbone using In- Fusion^®^ HD Cloning Kit (Takara). The WT donor template links rs7984-G in *cis,* and the mutant donor template links to rs7984-A in *cis* with the pathogenic autosomal dominant P347L mutation. The WT or mutant double-stranded donor plasmid was then co-injected with CRISPR/Cas9-gRNAs ribonucleoprotein (RNP) via microinjection into mouse zygotes. After implantation into pseudopregnant mice, the resulting F_0_ founders were then genotyped using primers in **Supplementary Table 2**. The founders carrying knock-in positive genotype results were backcrossed to C57BL/6J mice for 3 generations and fully sequenced. The respective homozygous WT and P347L mouse line were generated and used to obtain the experimental heterozygous c.-26A/G *RHO*^P347L/WT^ mice.

### Immunocytochemical staining, imaging and analysis

Cells were seeded on Vitronectin-coated glass coverslips and cultured under standard conditions. Prior to staining, the medium was aspirated, and cells were gently washed with PBS. Fixation was performed with 4% paraformaldehyde for 15 min at room temperature with gentle agitation. Cells were permeabilized with PBS containing 0.3% Triton X-100 for 5 minutes, followed by blocking in PBS containing 0.1% Triton X-100 and 3% bovine serum albumin for 1 h at room temperature with slow agitation. Primary antibodies, diluted in blocking solution, were applied for 1 h at room temperature under constant agitation. Primary antibodies included mouse monoclonal anti-rhodopsin (RHO; 1:500; Thermo Scientific, Ret-P1 clone), anti-mCherry (Invitrogen). After washing with PBS, cells were incubated with fluorophore-conjugated secondary antibodies (1:500 dilution), diluted in blocking solution, for 1 h at room temperature. Nuclei were counterstained with 4’,6-diamidino-2-phenylindole (DAPI; 1 µg/mL) for 3 min, followed by two washes with PBS. Coverslips were mounted using Fluorescence Mounting Medium (DAKO, Agilent Technologies). Z-stack fluorescence images were acquired using an Axio Imager Z1 microscope equipped with an ApoTome unit (Carl Zeiss) and analyzed with ZEN 2 Blue Edition software (Carl Zeiss).

For immunocytochemistry, RO were cultured for 20 days, fixed in 4% PFA, cryosectioned and stained as previously described (59). Primary antibodies and chemicals: anti-SAG (Thermo Fisher Scientific, PA1-731), anti-Rhodopsin (RHO; 1:500; Thermo Scientific, Ret-P1 clone), Phalloidin 568 (Thermo Fisher Scientific). RO cryosections were imaged on a Zeiss Apotome with a 63× objective using exposure-matched settings. Analysis was performed in Python (NumPy, SciPy, scikit-image, Matplotlib) assisted by Claude Coding. 182 fields from 75 organoids were analysed. Each z-stack was reduced to a maximum-intensity projection (MIP). OLM was traced by hand in Fiji using phalloidin channel visualizing the OLM. The signed distance of every pixel to this line was computed (apical/segment side negative, inner/nuclear side positive). Rhodopsin signal was median-filtered (3 × 3) before intensities were read. Fluorescence amount was measured as the background-subtracted mean above a fixed threshold set once per marker and batch Apical rhodopsin was quantified both as the amount within the outside-OLM zone (approximately 1–26 µm apical) and as a profile in 5 µm shells spanning 25 µm apical to 50 µm inside the OLM. To account for batch-to-batch intensity differences, each field was normalised to its own batch WT control before pooling. Statistical testing was performed using a two-sided Mann–Whitney U test (R135W vs. WT and R135W RHOligo-A 15 µM vs. R135W). For the distance profiles the test was applied shell by shell.

### ASO screening in retinal organoids

HiPSC-derived ROs were differentiated based on a protocol by Zhong et al. (60) with some modifications. Briefly, for embryoid body (EB) formation, 2.88 × 106 hiPSCs were detached on day 0 using TrypLE (ThermoFisher Scientific) and dissociated to single cells. Cells were then mixed with PeproGrow (Peprotech) medium, 10 µM Y-27632 (ROCK-inhibitor, Ascent Scientific) and 10 µM blebbistatin (Sigma-Aldrich) and distributed to 96 untreated V-shaped 96-wells (Sarstedt). For re-aggregation, the plate was centrifuged at 400 g for 4 min. On day 1, 80% of the medium was removed and replaced with N2 medium (DMEM/F12 (1:1)+Glutamax supplement (ThermoFisher Scientific), 24 nM sodium selenite (Sigma-Aldrich, USA), 16 nM progesterone (Sigma-Aldrich), 80 µg/ml human holotransferrin (Serologicals), 20 µg/ml human recombinant insulin (Sigma-Aldrich), 88 µM putrescin (Sigma-Aldrich, USA), 1x minimum essential media-non essential amino acids (NEAA, ThermoFisher Scientific, USA), 1x antibiotics-antimycotics (AA, ThermoFisher Scientific, USA)). Medium was changed again on day 4. On day 7, EBs were plated on Growth-Factor-Reduced Matrigel (BD Biosciences, USA)-coated six well plates at a density of 32 EBs/well and medium was changed daily. On day 16, medium was switched to a B27-based Retinal differentiation medium (BRDM) (DMEM/F12 (3:1) with 2% B27 (w/o vitamin A, ThermoFisher Scientific, USA), 1x NEAA and 1x AA). On day 24, eye fields were detached using 10 µl tips and collected in 10 cm bacterial petri dishes (Greiner Bio One, Germany) with BRDM. After completed formation, ROs were selected and if necessary detached from non-retinal spheres using micro scissors. From day 40 onwards, ROs in BRDM were supplemented with 10% fetal bovine serum (FBS, Thermo Fisher Scientific) and 100 µM taurine (Sigma-Aldrich). From day 70– 120, RO were supplemented with 1 µM retinoic acid (Sigma-Aldrich).

For the ASO treatment, human-derived ROs were selected, and the medium was fully replaced with BRDM lacking FBS and then distributed individually into non-adherent U-bottom 96-well plates (100 µL per well). For treatment, 50 µL of medium was replaced with treatment media containing ASO to achieve final concentrations of 5 µM or 15 µM; corresponding control conditions included scrambled oligonucleotides (SCR) or vehicle-only medium. Organoids were maintained for 48 h under standard culture conditions (20% O₂, 5% CO₂), followed by regular medium changes every 2–3 days using BRDM supplemented with 10% FBS and 100µM Taurine. Organoids were then individually collected, washed with PBS, lysed in RLT buffer, and stored at −80 °C prior to RNA extraction.

### Screening in the RHO-mCherry reporter cell lines

ASOs were delivered into RHO–mCherry reporter HEK293T cells by lipofection. Briefly, 4 × 10⁵ cells were seeded per well of a 12-well plate in DMEM supplemented with 10% fetal bovine serum (FBS) and the appropriate selection antibiotic. The following day, the medium was replaced with 900µl serum- and antibiotic-free DMEM, and cells were transfected with 50 nM ASO using Lipofectamine® 3000 according to the manufacturer’s instructions, with the exception that per transfection only 0.6 µl Lipofectamine 3000 and no P3000 enhancer were used. For flow cytometry, cells were detached by trypsinization and resuspended in phosphate-buffered saline (PBS) at 1 × 10⁶ cells/mL. Non-transfected cells served as negative controls. Data were acquired on a BD FACS Fortessa flow cytometer at the Flow Cytometry Core Facility Berg, Medical Faculty of the University of Tübingen. Cells were gated based on forward and side scatter to exclude debris, and singlets were identified using FSC-A versus FSC-H. RHO–mCherry fluorescence was detected using a 561 nm laser. Data were analyzed using FlowJo™ software (BD Biosciences). RHO–mCherry protein levels were quantified as median mCherry fluorescence intensity and normalized to cells treated with a scramble ASO of identical length, expressed as a percentage of the scramble control.

### *In-silico* off-target nomination

To nominate potential off-targets, a bioinformatic pipeline was developed to identify genomic sequences with high homology to the ASO sequence. First, potential off-target sites were identified using the GGGenome Browser aligning the respective sequence against the human reference genome (GRCh38/hg38). Searches were conducted across three distinct genomic contexts: curated pre-spliced RefSeq transcripts, curated spliced RefSeq transcripts, and the complete GRCh38.p13 assembly. To prioritize the most likely off-target candidates, results were filtered for sequences containing at least 14 matching nucleotides (0-2 mismatches/bulges). For off-targets located in genomic regions (p13 assembly), the Mutalyzer 2 Position Converter was utilized to map genomic coordinates to specific transcripts. To assess the potential impact of these hits on RNA processing, the distance of the off-target site to the nearest splice junction was calculated based on HGVS nomenclature. To evaluate the biological relevance of the nominated off-targets, the pipeline integrated expression data from the Human Protein Atlas API.

## Animal experiments

Intravitreal injections were performed on anesthetized mice using glass microcapillary needles (FIVEphoton Biochemicals^TM^) attached to a manual microinjection pump. Briefly, the eye was gently protruded, and the injection was performed slightly below the level of the ciliary muscles at the limbus, inserting the needle at an approximately 45° angle to avoid lens damage. A volume of 2 µL ASO was injected into the vitreous humor. Animal breeding and experiments were approved and performed under IACUC regulations.

## Electroretinography

Electroretinography (ERG) was performed as previously described (61). Briefly, animals were dark-adapted overnight and anesthetized with ketamine (100mg/kg) xylazine (10mg/kg) injected intraperitoneally. Pupils were dilated using 0.5% tropicamide. Scotopic (Rod) and photopic (Maximum) ERG responses were recorded following stimulation with pulsed white light flashes (6,500 K) at intensity of 0.00130 cd/m^2^ and 3 cd/m^2^. After a 10-min light adaptation, the photopic (Cone) response was recorded with pulsed flashs at intensity of 30 cd·s/m^2^. ERG results were collected and analyzed using Espion Software V6.67.10 (Diagnosis LLC).

## Retina histology

Mice were euthanized by anesthetic overdosing, and eyes were enucleated, fixed, and paraffin-embedded. Retinal cross-sections (10 µm) were stained with hematoxylin and eosin (H&E) as previously described (61). Images were acquired using a Leica SCN400 whole-slide digital imaging system and analyzed with Aperio ImageScope software (Leica).

## Quantitative reverse transcription PCR (qRT-PCR) and allele-specific assay

To quantify gene expression levels, total RNA was isolated from HEK293T cells, retinal organoids, and mouse retinae using the GenUP™ Total RNA Kit (Biotechrabbit), RNeasy Micro Kit (Quiagen) and RNeasy MinElute Cleanup Kit (Qiagen), respectively. Depending on experiment, 50 to 500 ng of total RNA was treated with DNase I (Invitrogen) to remove genomic DNA contamination and subsequently reverse-transcribed into complementary DNA (cDNA) using the OneScript® Plus cDNA Synthesis Kit (abm), the SuperScript IV First Strand Synthesis Kit (Thermo Fisher Scientific) or the LunaScript® RT SuperMix Kit (New England Biolabs, NEB) respectively, according to the manufacturer’s instructions. Quantitative PCR (qPCR) was performed using Power SYBR™ Green PCR Master Mix (Thermo Fisher Scientific) for HEK293T samples and TaqMan™ Fast Advanced Master Mix (Thermo Fisher Scientific) for mouse retinae. Primer sequences and TaqMan probe details are provided in **Supplementary Table 2**. For the determination of the *RHO* to *GUSB* expression ratio in post- mortem human retinae, RNA sequencing data from Pinelli et al., 2016 were used (62). The number of reads corresponding to *RHO* transcripts was divided by the number of *GUSB* transcripts in the 19 highest-quality samples of the dataset. Relative expression levels were calculated using the ΔΔCt method after normalization to housekeeping genes.

## Immuno blot

Retinal tissue was isolated from enucleated mouse eyes and homogenized in Mem-PER™ lysis buffer (Thermo Fisher Scientific) supplemented with protease and phosphatase inhibitors (Sigma-Aldrich, P2714). Tissue dissociation was performed under constant agitation for 30 min at 4 °C, followed by sonication. Lysates were cleared by centrifugation at 20,000 × g for 10 min at 4 °C. Protein concentrations were determined by Bradford assay (Bio-Rad), and 3 µg of total protein per sample were mixed with 4× Laemmli buffer (Bio-Rad) and incubated for 60 s at room temperature. Proteins were separated using Mini-PROTEAN® TGX™ precast gels (Bio-Rad) and transferred onto polyvinylidene difluoride (PVDF) membranes.

Membranes were blocked for 30 min in EveryBlot™ blocking buffer (Bio-Rad) and incubated overnight at 4 °C with primary antibodies diluted in blocking buffer. Primary antibodies included mouse monoclonal anti-rhodopsin (RHO; 1:5000; Thermo Fisher Scientific, 1D4 clone) and rabbit polyclonal anti-recoverin (RCVN; 1:500; Sigma Aldrich, AB5585-I). Following three washes with PBST (5 min each), membranes were incubated for 1 h at room temperature with Horseradish peroxidase (HRP)-conjugated secondary antibodies: rabbit anti-mouse IgG (1:20,000; Abcam, ab6728) or mouse anti-rabbit IgG (1:20,000; Abcam, ab99697). After additional PBST washes, immunoreactive bands were detected using Immobilon Western chemiluminescent substrate (Millipore, WBKLS0500) and visualized using a Med-Dent developer/fixer system (Z&Z Medical, 2500DF).

### RNA-sequencing

Total RNA extraction was performed using the RNeasy Micro Kit (QIAGEN). RNA-sequencing was performed by Novogene GmbH using poly-A enrichment protocol and sequencing on NovaSeq X Plus Series (PE150) (6 G raw data per sample)(63). Primary reads alignment analysis was performed using HISAT2 by Novogene GmbH on their online platform NovoMagic .

Total RNA extraction was performed using the RNeasy Micro Kit (QIAGEN). RNA-sequencing was performed by Novogene GmbH using poly-A enrichment protocol and sequencing on NovaSeq X Plus Series (PE150) (6 G raw data per sample). Primary reads alignment analysis was performed using HISAT2 by Novogene GmbH on their online platform NovoMagic. Downstream analyses were performed in R (version 4.4.2). Genes with fewer than 10 total counts across all samples were excluded from analysis. Differential expression analysis was performed using DESeq2 (version 1.44) (64) with a Wald test and a negative binomial generalized linear model. For retinal organoid samples (n = 3 per group: NTC, SCR, RHOligo- A), the design formula ∼condition was used and three pairwise contrasts were computed: RHOligo-A vs. NTC, RHOligo-A vs. SCR, and SCR vs. NTC. For RPE monolayer samples (n = 5 per group: NTC, SCR, SILE, RHOligo-A, P23H, BAD), pairwise contrasts versus NTC were computed using the same framework. Genes with a Benjamini-Hochberg adjusted p-value < 0.05 and |log2 fold-change| > 1 were classified as differentially expressed (DEGs). To account for variable RPE cell contamination between different RO samples, a per-sample RPE contamination score was calculated as the mean z-score of ten established RPE marker genes (RPE65, BEST1, RLBP1, RGR, TYRP1, DCT, PMEL, MITF, TTR, CLDN10) across variance-stabilized count data (DESeq2 vst). This score was included as a continuous covariate in a secondary DESeq2 model (∼rpe_score + condition). DEGs that lost significance after this correction were considered confounded by RPE contamination rather than driven by RHOligo-A sequence specificity.

DEG lists from each contrast were intersected with the predicted off-target candidate list (n = r length(ot_genes) genes). Results are visualized as a four-set Venn diagram integrating the in silico prediction set, organoid DEGs from the naive and RPE-corrected models, and RPE cell DEGs. Statistical analysis, figure generation, and data export were performed using the R packages DESeq2, ggplot2 (65), patchwork, and ggvenn. All code and intermediate result tables are available upon request.

### Transmission electron microscopy

Mice were fixed by cardiac perfusion in 2% Paraformaldehyde (PFA), 2,5% Glutaraldehyde (GA) in 0,1 M sodium cacodylate buffer. After dissection, eyebulbs were postfixed in 4% PFA, 2,5% GA in cacodylate buffer for 2 days.

After fixation the samples were rinsed three times in 0.1 M sodium cacodylate buffer (pH 7.4) for a total of 30 minutes, and postfixed in 1 % OsO4 for 1.5 hours at room temperature (Electron Microscopy Sciences). After three additional washes in cacodylate buffer and dehydration in 30 % and 50 % ethanol, the tissue was counterstained with 3 % uranyl acetate dissolved in 70 % ethanol (Serva) overnight, followed by graded ethanol concentrations up to 100 % (15 min steps, 1 x 80% - 96% and 3 x 100%, propylene oxide 2 x 15 min). The dehydrated samples were incubated in a 2:1, 1:1 and 1:2 mixture of propylene oxide and Epon resin (Serva) for 1 hour each. Finally, samples were infiltrated with pure Epon for 2 hours. Samples were embedded in fresh resin in block molds and cured for 3 days at 60°C.

Ultrathin sections (50 nm) were cut on a Reichert Ultracut S (Leica), collected on copper grids and counterstained with Reynolds lead citrate. Sections were analyzed with a Zeiss EM 900 transmission electron microscope (Zeiss) equipped with a 2k x 2k CCD camera.

### PBMC cytokine release assay

In order to establish the initial immunostimulatory profile of RHOligo_A (A-ASO-5) in *in vitro* settings, peripheral blood mononuclear cells (PBMCs), isolated from buffy coats of nine healthy donors upon written informed consent, were exposed to RHOligo_A for 24 h at 37 °C with 5% CO2 at final concentrations of 1 μM and 10 μM. Control ASO, an ASO targeting RHO:p.P23H, previously investigated (37), and PBS were included as negative controls. Resiquimod (R848; Invivogen, tlrl-r848-1) and polyinosinic:polycytidylic acid [poly(I:C); Invivogen, tlrl-pic] served as positive controls as TLR7/8 and TRL3 agonists, respectively. After 24 h, culture supernatants were collected and the concentrations of IL-6 (R&D systems DY206-5), TNF-α (R&D systems DY210-5), IFN-α (MABTECH 3425-1H-6), IP-10 (R&D systems DY266-5), MIP-1α (R&D systems DY270-5) and MIP-1β (R&D systems DY271-5) were determined by enzyme-linked immunosorbent assays (ELISA) according to the manufacturers’ instructions. Proinflammatory marker concentrations were determined from standard curves and analyzed relative to PBS-treated samples for each donor.

## Statistical analysis

Data was analyzed using GraphPad Prism software. Statistical tests and significance thresholds are specified in the figure legends.

## ACKNOWLEDGMENTS

We thank Dr. Martin Maier, for his support and expert advice, Dr. Neoklis Makrides for performing paraformaldehyde fixation of mice, members of Jonas Children’s Vision Care (JCVC) for sharing mice and ideas, and Prof. Dr. Mihai Netea and Andrei Sarlea for facilitating the PBMC assay. PDA is supported by Tistou and Charlotte Kerstan Foundation and Hector Fellow Academy. JCVC is supported by the National Institute of Health U01 EY030580, U01EY034590 R24EY028758, R24EY027285, 5P30EY019007, R01EY033770. R01EY018213, R01EY024698, the Foundation Fighting Blindness TA-GT- 0321-0802-COLU-TRAP, Richard Jaffe, NYEE Foundation, the Rosenbaum Family Foundation, Schneeweiss Stem Cell Fund, the Gebroe Family Foundation, Piyada Phanaphat fund, the Research to Prevent Blindness (RPB) Physician-Scientist Award, unrestricted funds from RPB, New York, NY, USA.

---

**Supplementary Figure 1.**
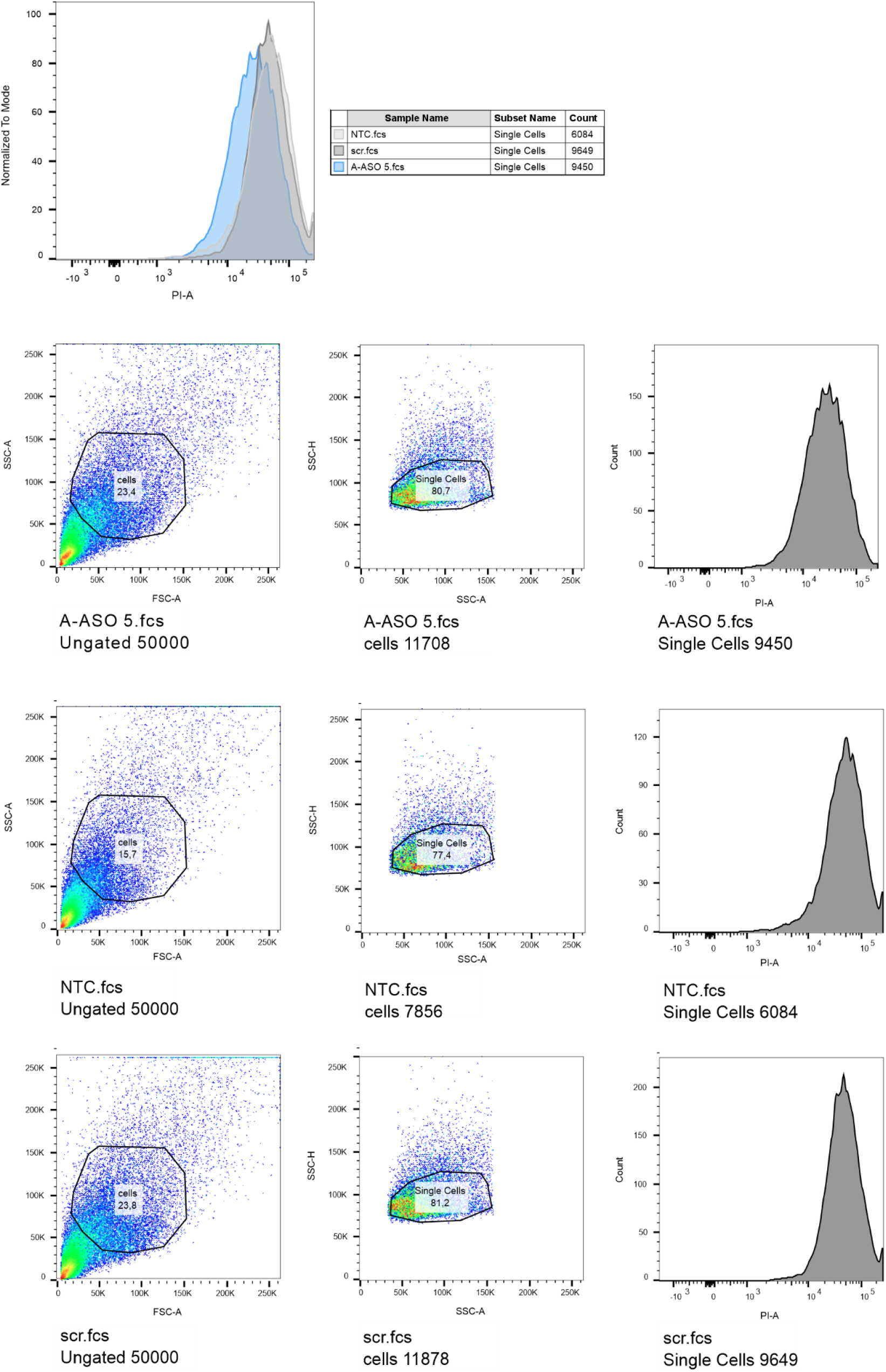
**Example for flow cytometry for A-ASO screening**Overlay flow cytometry histogram (top) and separate scatter blots and histograms (bottom) of a representative replicate of RHO:c.-26A+c.696+4T reporter cells transfected with A_ASO5 and scramble (scr) ASO compared to non-transfected control (NTC). PI channel used to detect mCherry fluorescence.

**Supplementary Figure 2.**
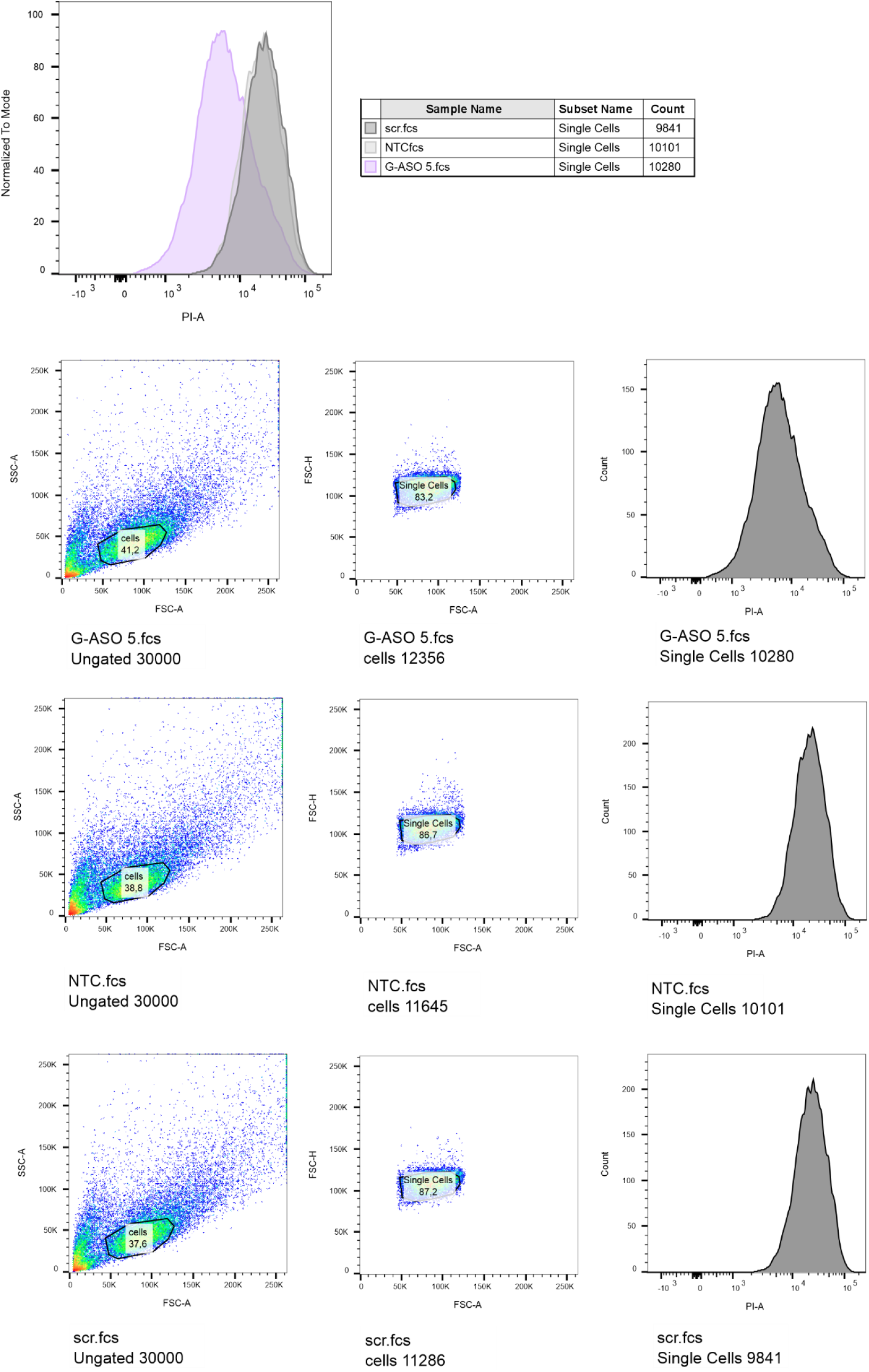
**Example for flow cytometry for G-ASO screening**Overlay flow cytometry histogram (top) and separate scatter blots and histograms (bottom) of a representative replicate of RHOc.-26G+696+4C reporter cells transfected with A_ASO5 and scramble (scr) ASO compared to non-transfected control (NTC). PI channel used to detect mCherry fluorescence.

**Supplementary Figure 3.**
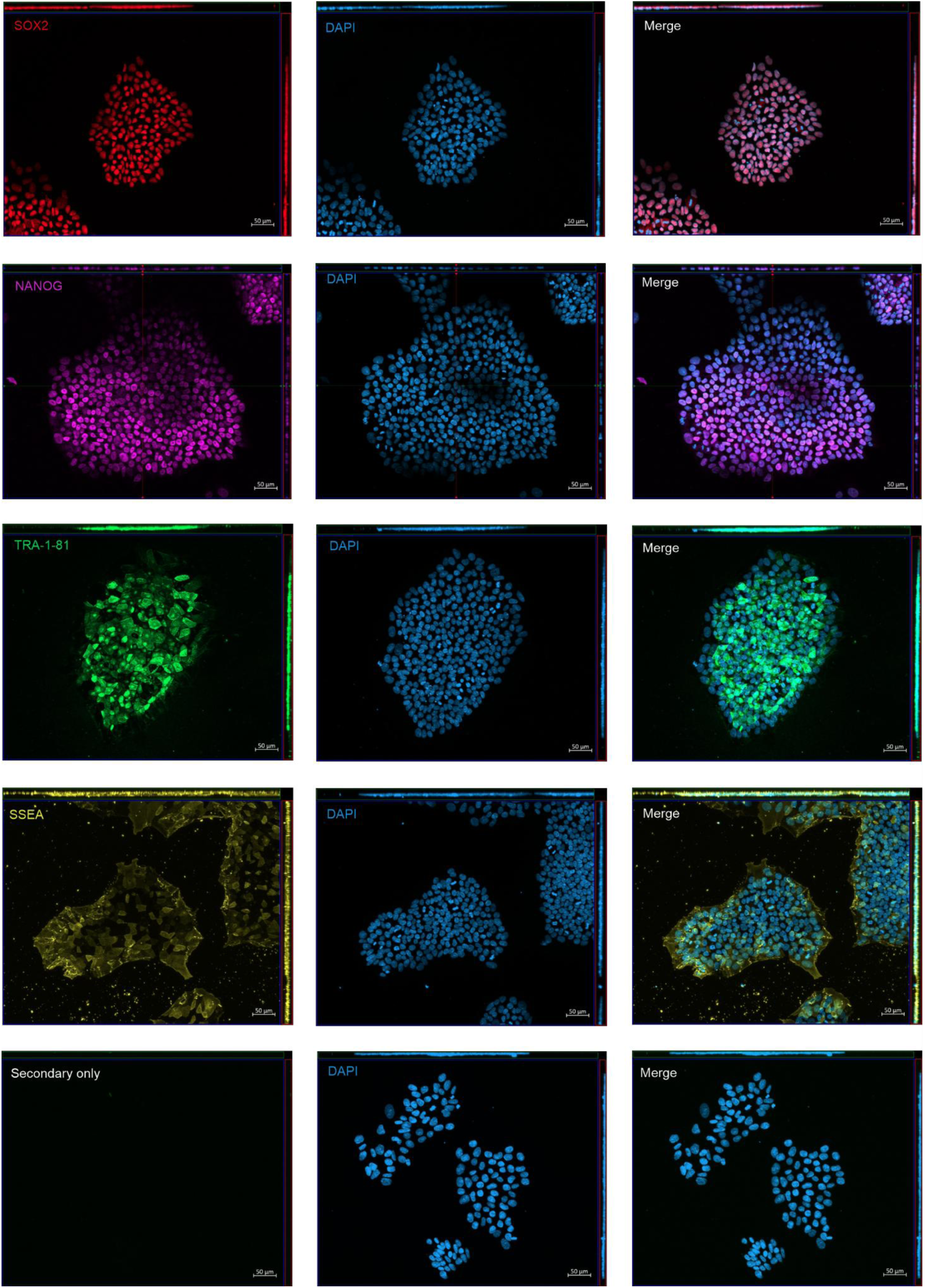
Pluripotency marker expression in c.-26A/G RHO^R135W/WT^ patient-derived iPSCs. Immunofluorescence staining demonstrates expression of established pluripotency markers in c.-26A/G RHO^R135W/WT^ patient-derived iPSCs. Nuclear markers SOX2 (red) and NANOG (magenta) are co-expressed with the cell surface markers TRA-1-81 (green) and SSEA (yellow). Nuclei are counterstained with DAPI (blue). A secondary antibody–only control was included to confirm staining specificity.

**Supplementary Figure 4.**
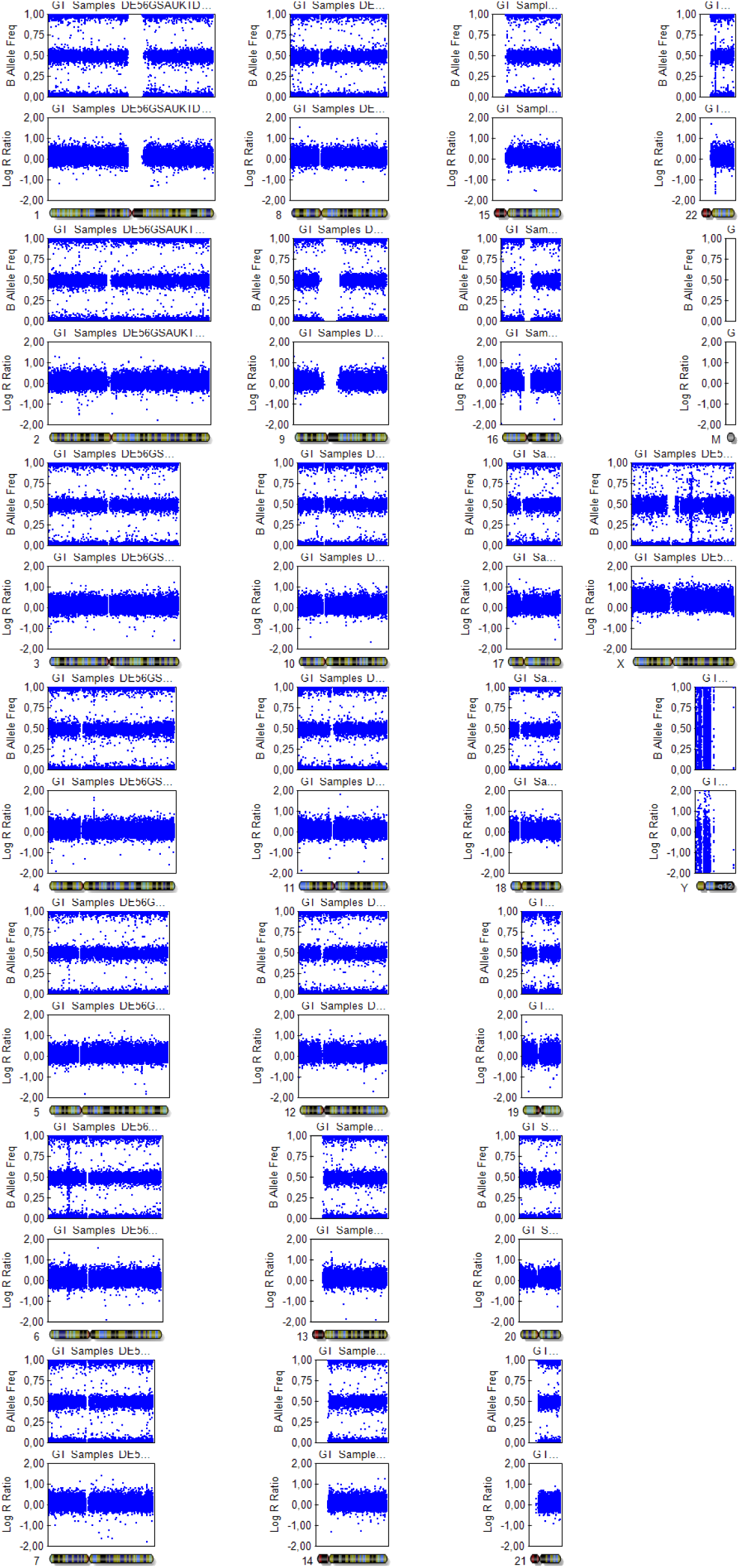
Copy number variation analysis of patient-derived c.-26A/G c.-26A/G *RHO*^R135W/WT^ iPSC clone. Genome-wide copy number variation (CNV) analysis was performed to assess the genomic integrity of the RHO mutant iPSC clone used in this study. B-allele frequency and log R ratio profiles are shown across all chromosomes. No major chromosomal abnormalities or copy number alterations were detected, indicating preservation of genomic stability.

**Supplementary Figure 5.**
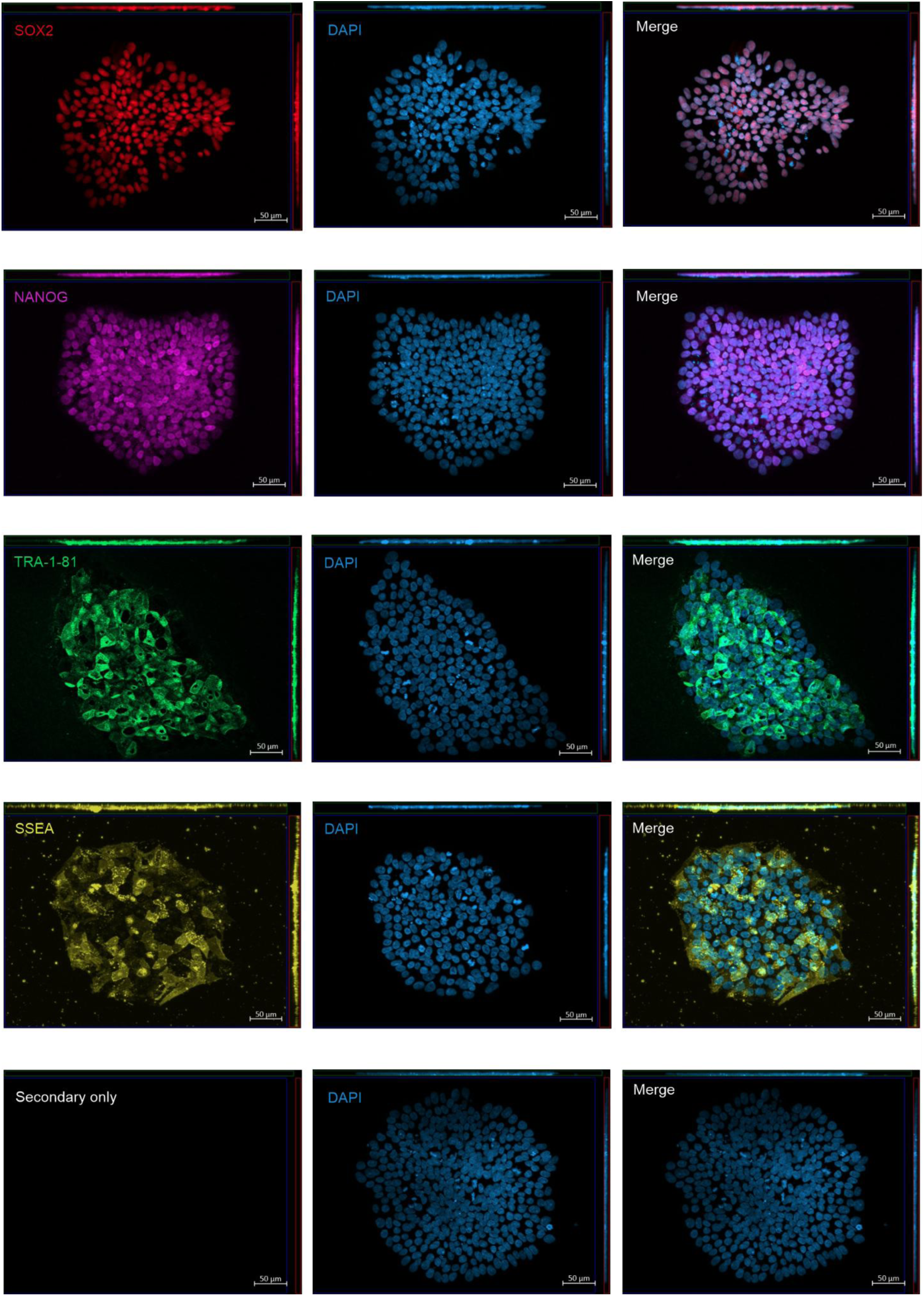
Pluripotency marker expression in c.-26A/G RHO^WT/WT^ patient-derived isogenic WT iPSCs. Immunofluorescence staining demonstrates expression of established pluripotency markers in c.-26A/G RHO^WT/WT^ patient-derived isogenic WT iPSCs. Nuclear markers SOX2 (red) and NANOG (magenta) are co-expressed with the cell surface markers TRA-1-81 (green) and SSEA (yellow). Nuclei are counterstained with DAPI (blue). A secondary antibody–only control was included to confirm staining specificity.

**Supplementary Figure 6.**
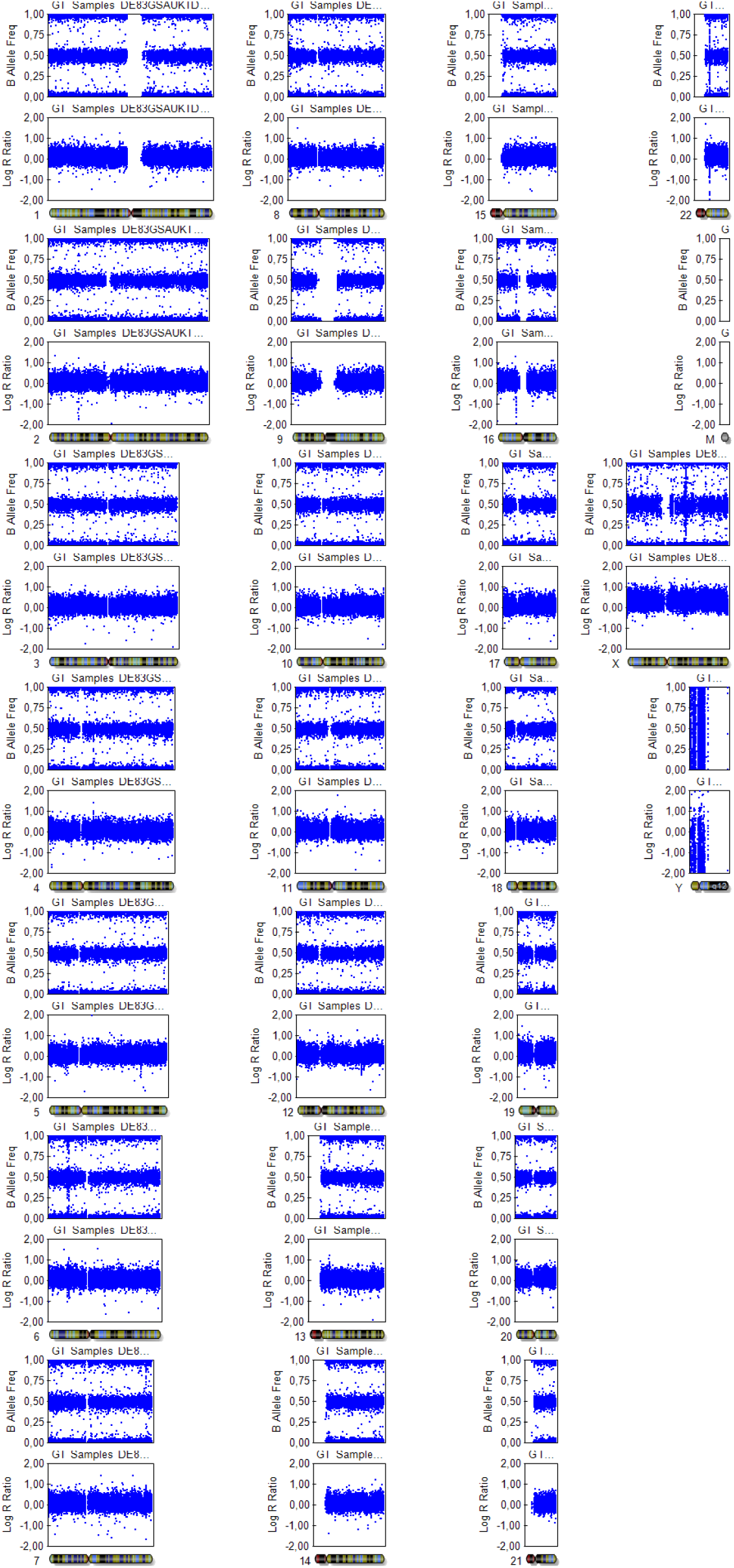
Copy number variation analysis of the isogenic wild-type c.- 26A/G c.-26A/G *RHO*^WT/WT^ iPSC clone. Genome-wide copy number variation (CNV) analysis was performed to assess the genomic integrity of the isogenic wild-type iPSC clone used in this study. B-allele frequency and log R ratio profiles are shown across all chromosomes. No major chromosomal abnormalities or copy number alterations were detected, indicating preservation of genomic stability following CRISPR-mediated knock-in to correct c.403C>T p.R135W back to WT.

**Supplementary Figure 7.**
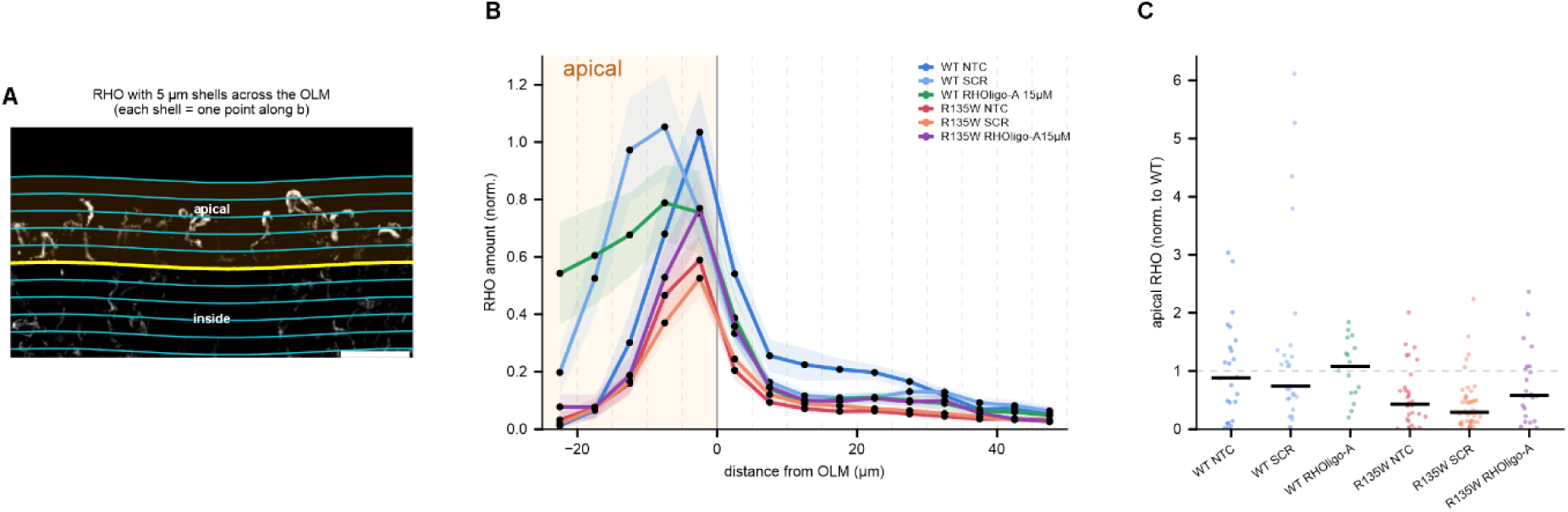
Rhodopsin distribution across the OLM for each WT and R135W mutant retinal organoid condition. (A) Schematic explanation the outer-limiting membrane (OLM) based shell separation (each shell ∼5µm) (B) Each WT and R135W is either non-transfected control (NTC), treated with scrambled (SCR) or 15 uM RHOligo-A. RHO amount versus distance from the OLM, in 5 µm shells from 25 µm apical (-) to 50 µm inside (+); dotted lines mark shell edges; orange area: apical side. Lines are mean ± SEM. across images, each normalised to its batch-WT peak shell. (C) Apical Rhodopsin amount (∼1-26 um zone apical to the OLM) per image, normalised to the batch-WT median. Signal was quantified with a fixed per-batch threshold.

**Supplementary Figure 8.**
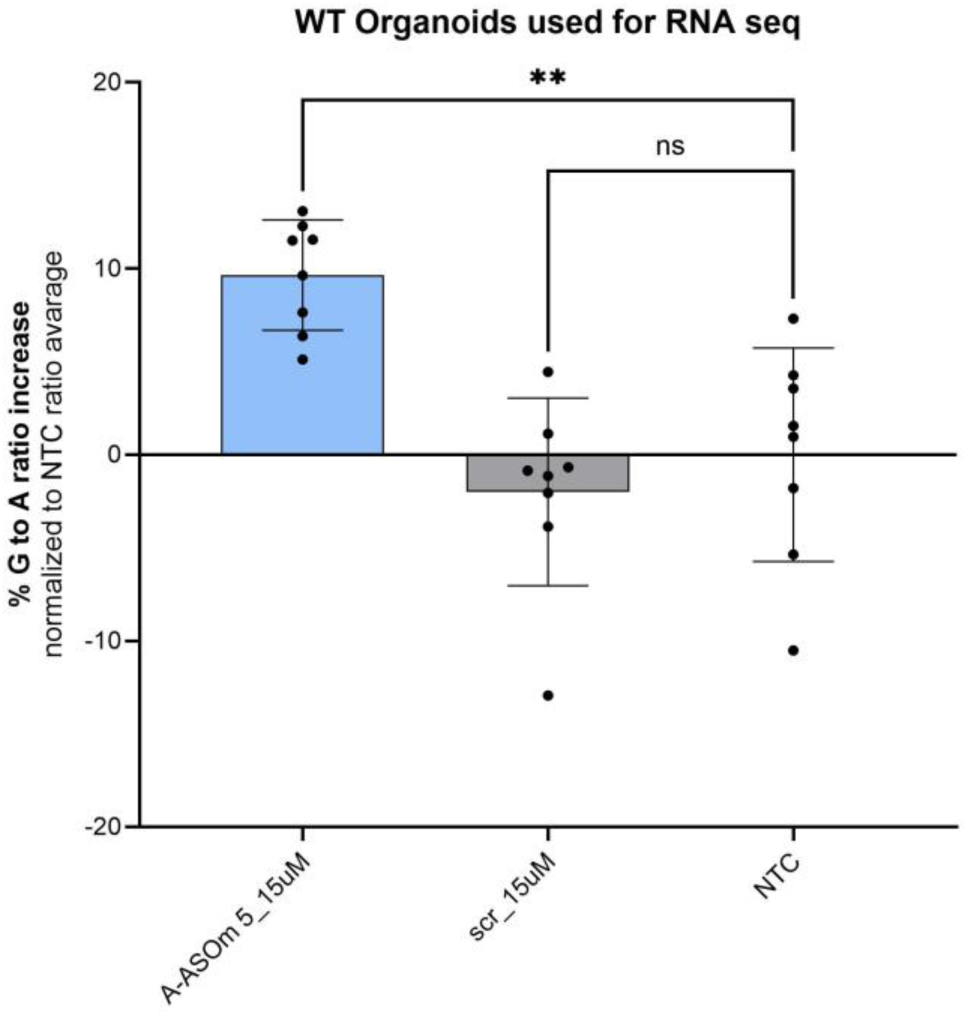
Relative *RHO* transcript abundance after ASO-treatment in c.- 26A/G RHO^WT/WT^ retinal organoids used for RNA-seq. iPSC-derived c.-26A/G *RHO*^WT/WT^ retinal organoids (n=8) were treated with 15µM RHOligo-A by gymnotic uptake and 10 days washout regiment. Next generation sequencing of cDNA determined change of relative decrease of A-allele when compared to both untreated (NTC) and scr-treated organoids. Statistical significance was determined using ordinary one-way ANOVA.**p ≤ 0.01. Data are shown as average ± standard deviation. Individual dots represent individual Retinal organoids.

**Supplementary Figure 9.**
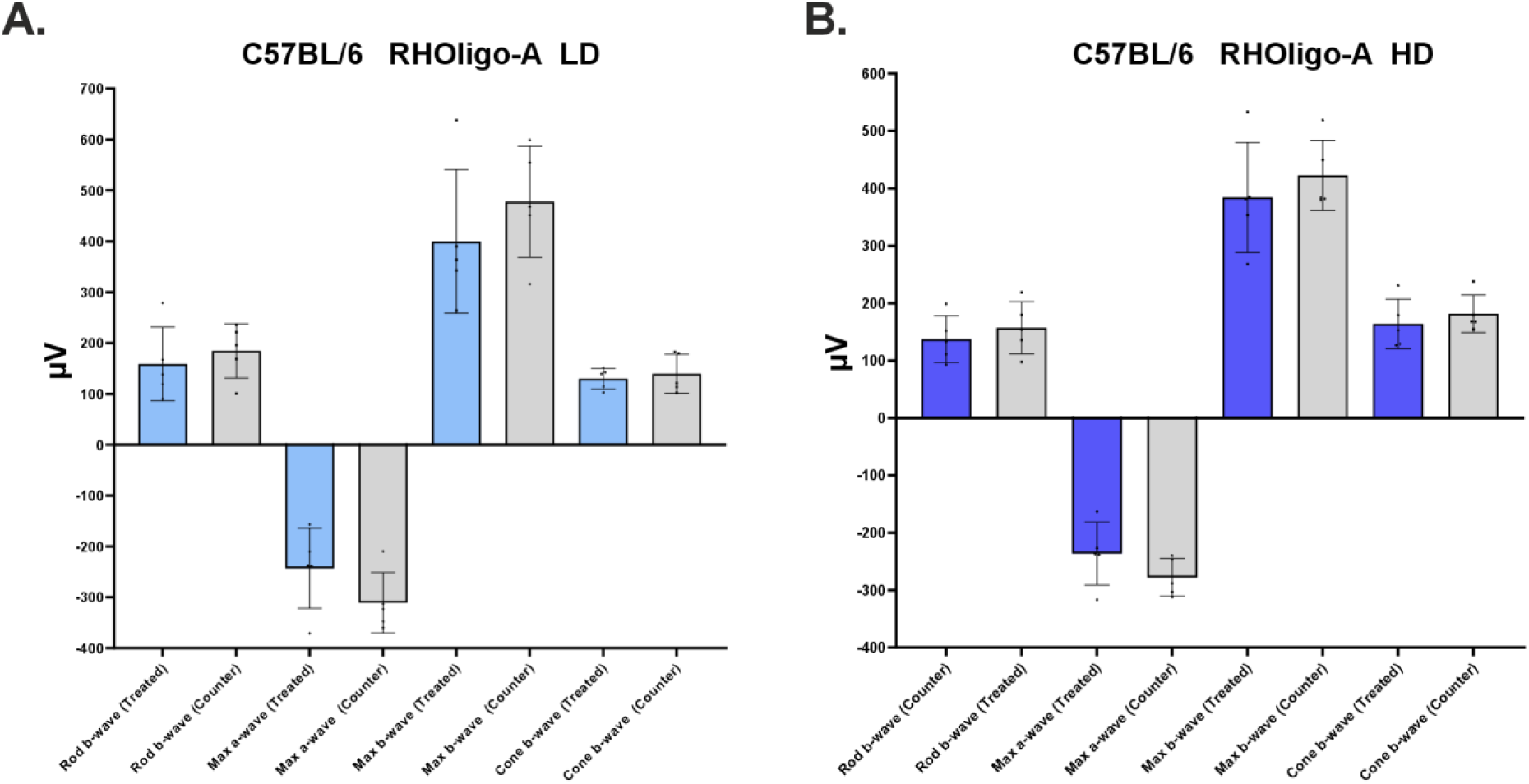
Ocular tolerability assessment of RHOligo-A in C57BL/6 mice. Six-week-old C57BL/6 mice received unilateral intravitreal injections of RHOligo-A to evaluate ocular tolerability at two doses: (**A**) low dose (LD, 10 µg) and (**B**) high dose (HD, 50 µg). Retinal function was assessed by electroretinography (ERG) 30 days post-injection. No statistically significant differences were observed between treated and contralateral control eyes across all ERG parameters at either dose, indicating a favorable ocular safety profile of the lead ASO candidate. Data are shown as average ± SEM. Individual dots represent individual values from different animals.

**Supplementary Figure 10.**
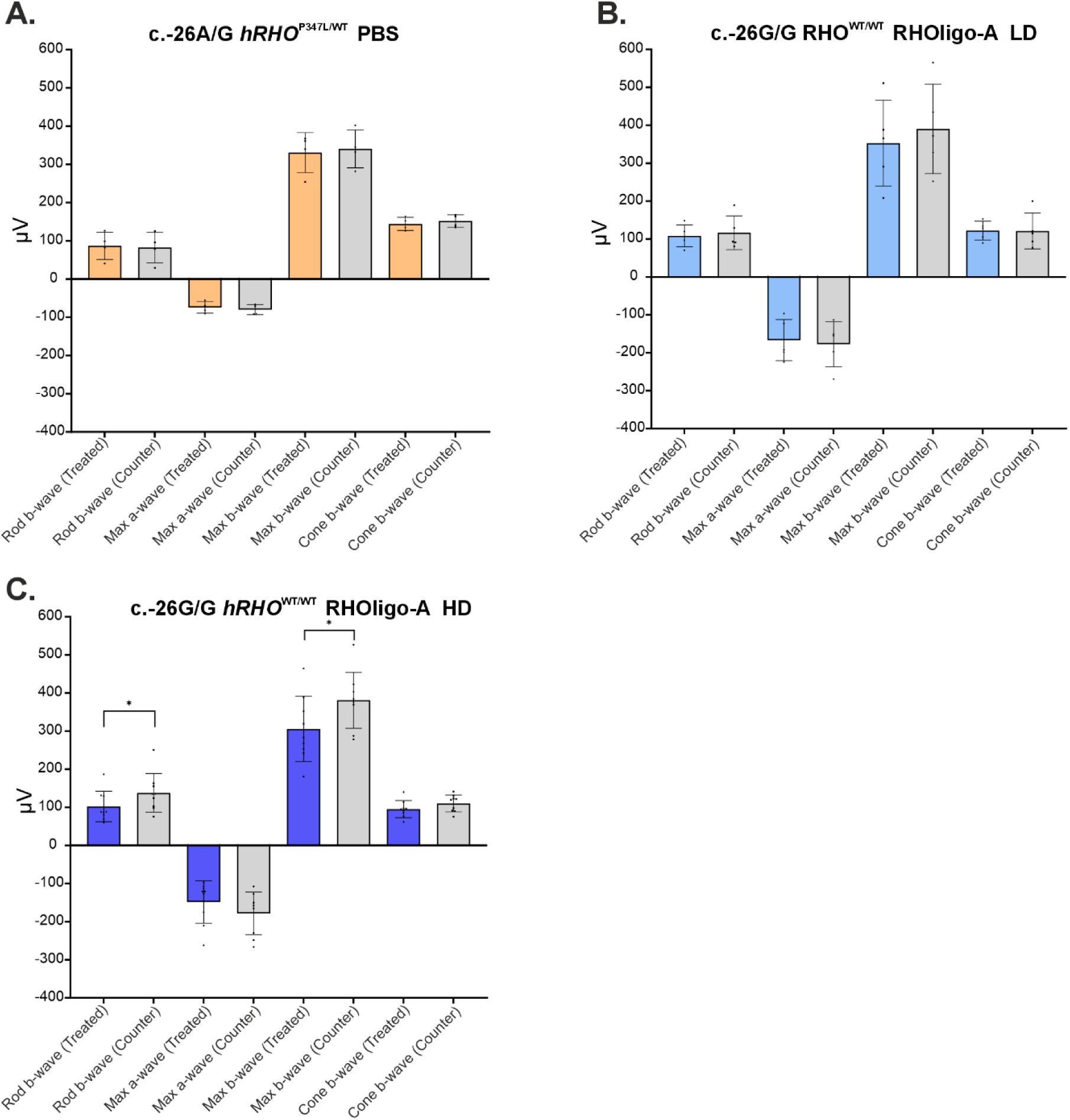
ERG responses of controls for intravitreal injection in c.- 26A/G h*RHO*^P347L/WT^ mice and effects of RHOligo-A on ERG responses in c.-26G/G h*RHO*^WT/WT^ mice. (A) To assess the potential impact of the intravitreal injection procedure on ERG responses, c.-26A/G h*RHO*^P347L/WT^ mice were unilaterally injected with PBS. Electroretinography (ERG) performed 30 days post-injection revealed no differences between injected and contralateral eyes, indicating that the procedure itself does not affect retinal function. (B–C) To evaluate potential effects of partial off-target RHO knockdown in wild-type animals, c.-26G/G h*RHO*^WT/WT^ mice were treated with RHOligo-A at low dose (LD, 10 µg) or high dose (HD, 50 µg). LD-treated mice showed no differences between treated and contralateral eyes across ERG parameters. In contrast, HD-treated mice exhibited a modest reduction in scotopic responses, including maximal b-wave amplitudes, while cone-specific responses remained largely unchanged. These findings suggest that partial *RHO* knockdown in wild-type retina may result in subtle functional effects at higher doses. Data are shown as average ± SEM. Individual dots represent individual values from different animals.

**Supplementary Figure 11.**
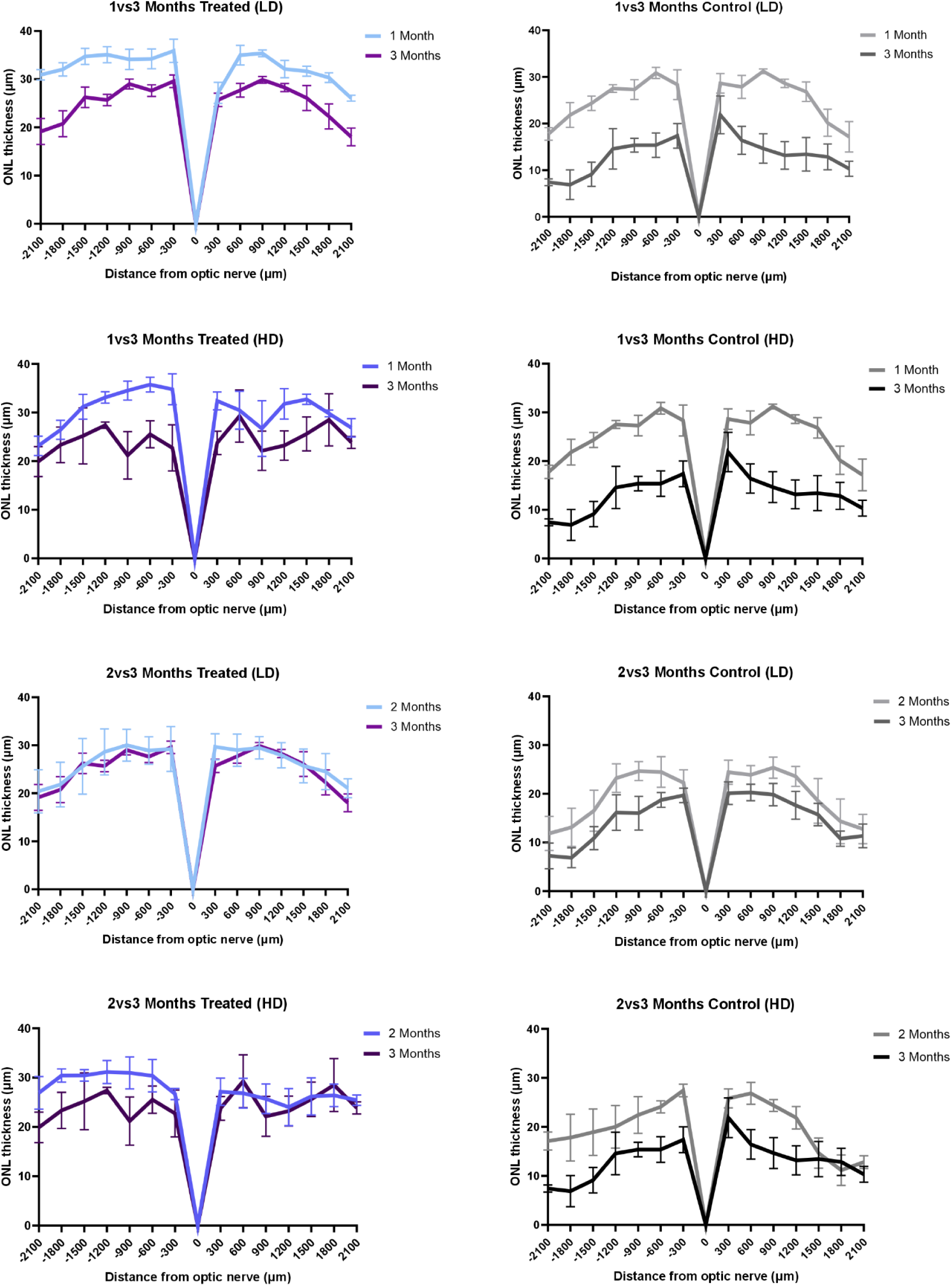
Comparison of retinal structural preservation over time. Spider plots illustrating outer nuclear layer (ONL) thickness across the retina are shown for comparisons between 1 and 3 months post-treatment, and between 2 and 3 months post-treatment, for both low-dose (LD) and high-dose (HD) groups. Corresponding contralateral untreated eyes are included for comparison. Data are presented as mean ± SEM.

**Supplementary Figure 12.**
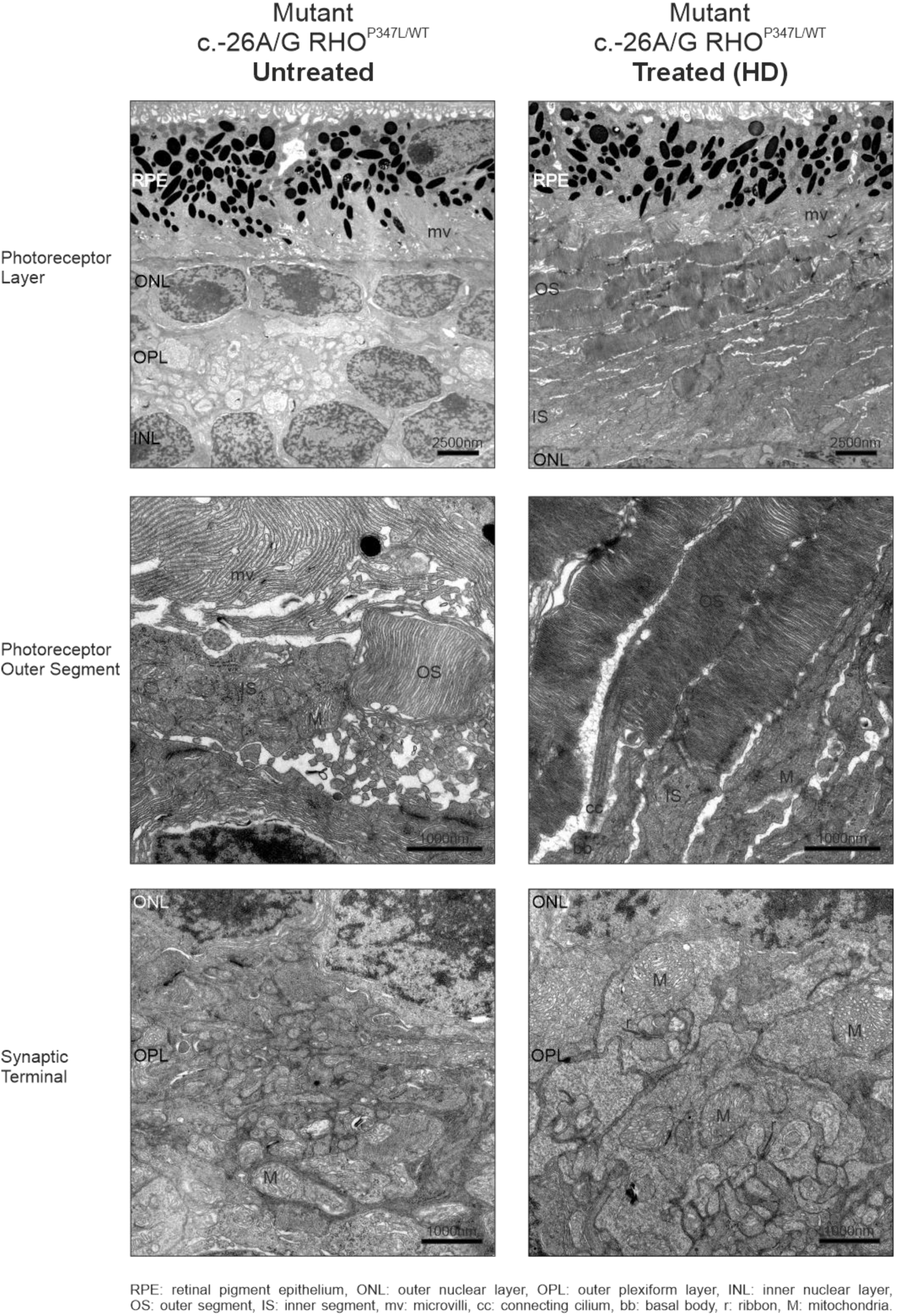
Ultrastructural analysis of retinal morphology by electron microscopy in RHOligo-A treated and untreated mutant c.-26A/G h*RHO*^P347L/WT^ mice. Electron micrographs of peripheral retina from high-dose (HD)–treated and untreated mice at 3 months post-injection. Photoreceptor layer (×3,000): Untreated mice show complete loss of photoreceptor outer segments, with only a residual layer of photoreceptor nuclei remaining. In contrast, HD-treated animals display preserved photoreceptor structure, including clearly defined and organized outer segments. Photoreceptor outer segments (×12,000): At higher magnification, untreated mice exhibit only sparse and disorganized outer segment remnants. Conversely, treated animals show well-structured outer segments with morphologically intact and properly stacked disc membranes. Synaptic terminals (x12,000): In the outer plexiform layer, untreated mice rarely exhibit functional synapses, as indicated by the near absence of synaptic ribbons. In contrast, treated animals display well-organized synaptic architecture, including prominent ribbon synapses and associated mitochondria, consistent with functional photoreceptor synaptic terminals.

**Supplementary Figure 13.**
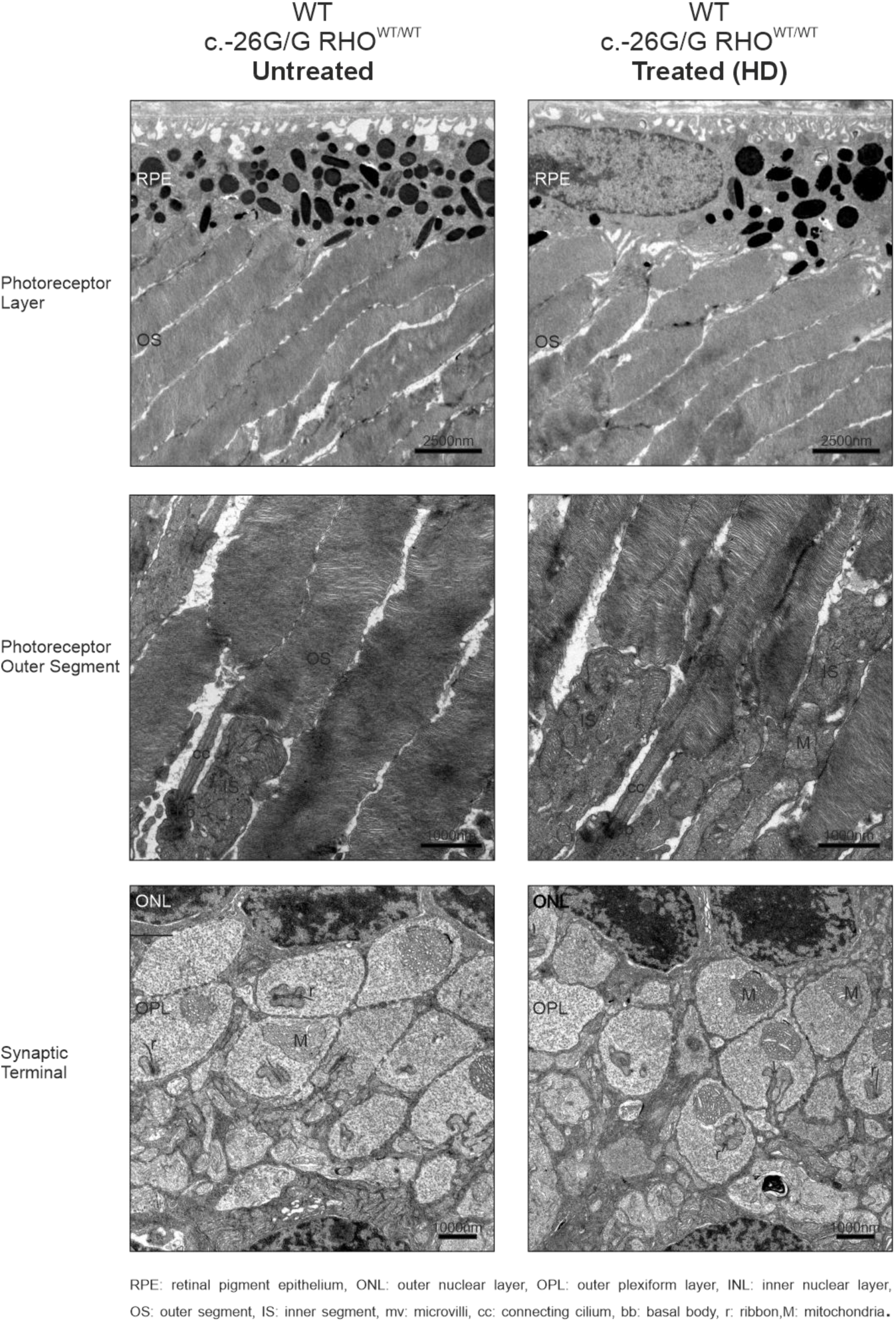
Ultrastructural analysis of retinal morphology by electron microscopy in RHOligo-A treated and untreated wild-type c.-26G/G h*RHO*^WT/WT^ mice. Electron micrographs of peripheral retina from high-dose (HD)–treated and untreated mice at 3 months post-injection. Photoreceptor layer (×3,000): Both untreated and HD-treated mice display comparable photoreceptor morphology, with preserved outer segments. However, outer segments appear tilted relative to the retinal pigment epithelium (RPE), rather than exhibiting the typical perpendicular (90°) orientation, suggesting a structural feature of this humanized model. Photoreceptor outer segments (×12,000): At higher magnification, untreated and treated mice exhibit similar ultrastructural organization, with morphologically intact and properly stacked disc membranes. Synaptic terminals (×12,000): In the outer plexiform layer, both untreated and treated mice show well-organized and functional synapses, as evidenced by clearly identifiable ribbon synapses and associated mitochondria, with no apparent differences between groups.

**Supplementary Table 1.**
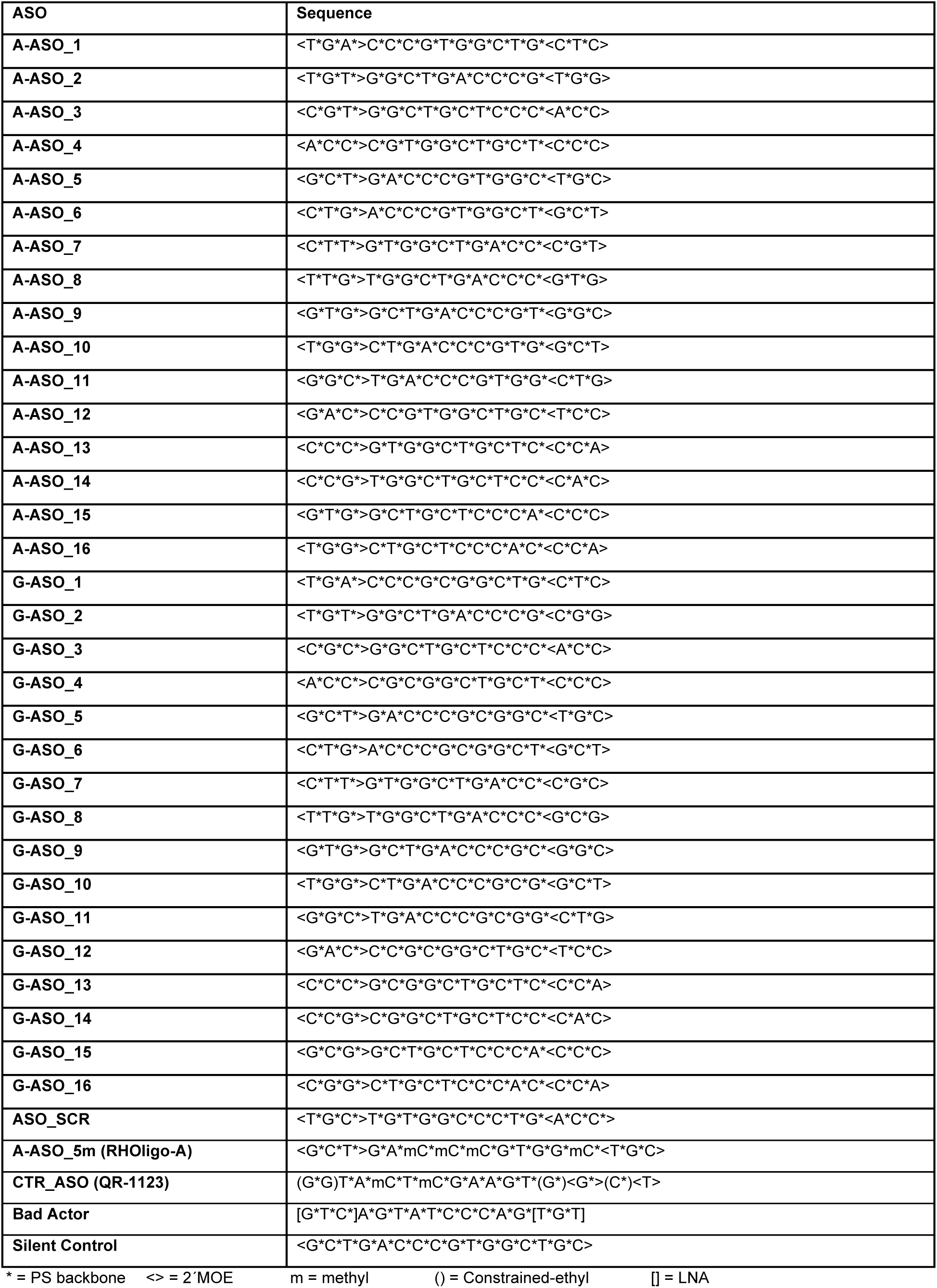
ASO Sequences

**Supplementary Table 2.**
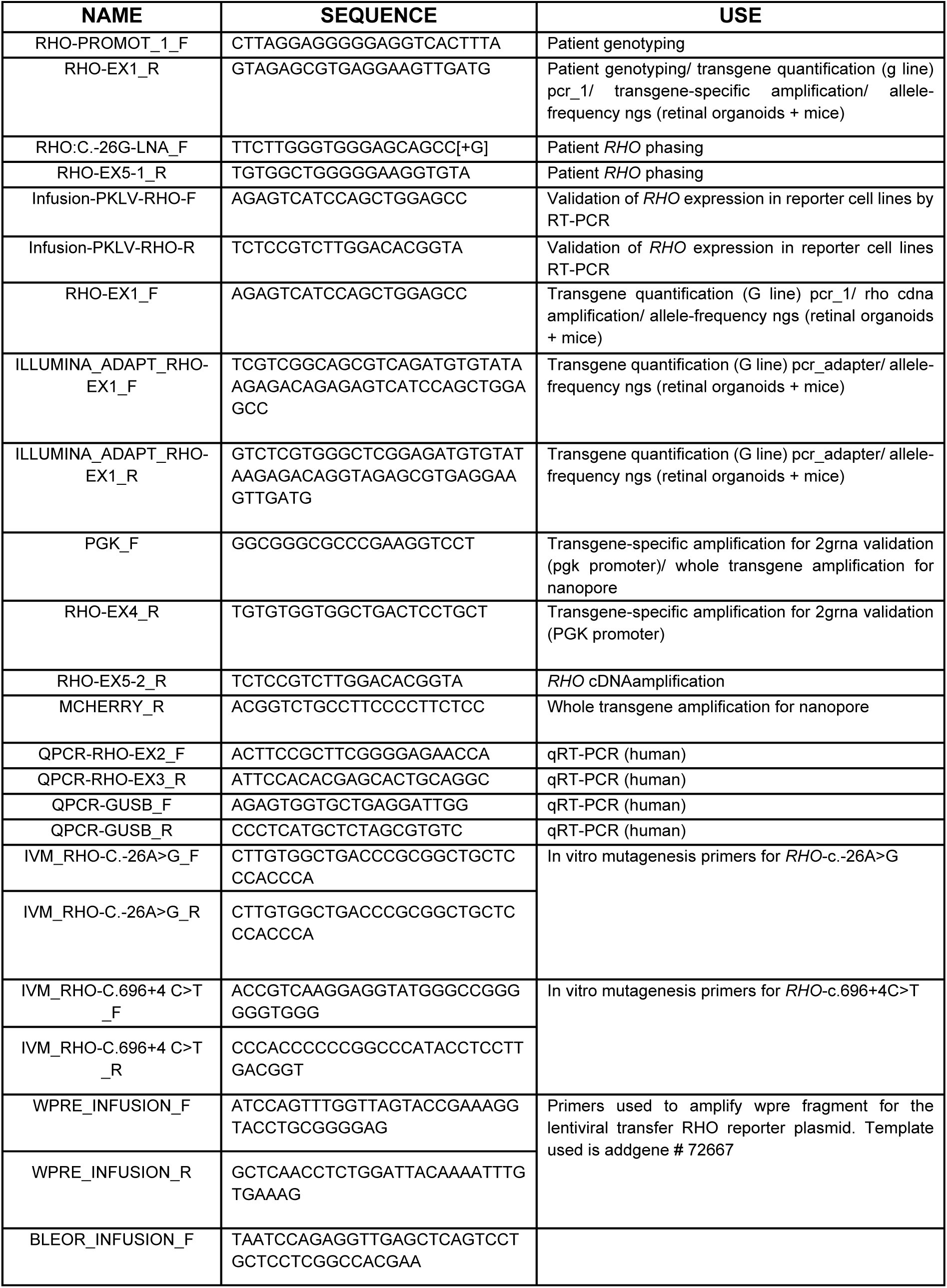
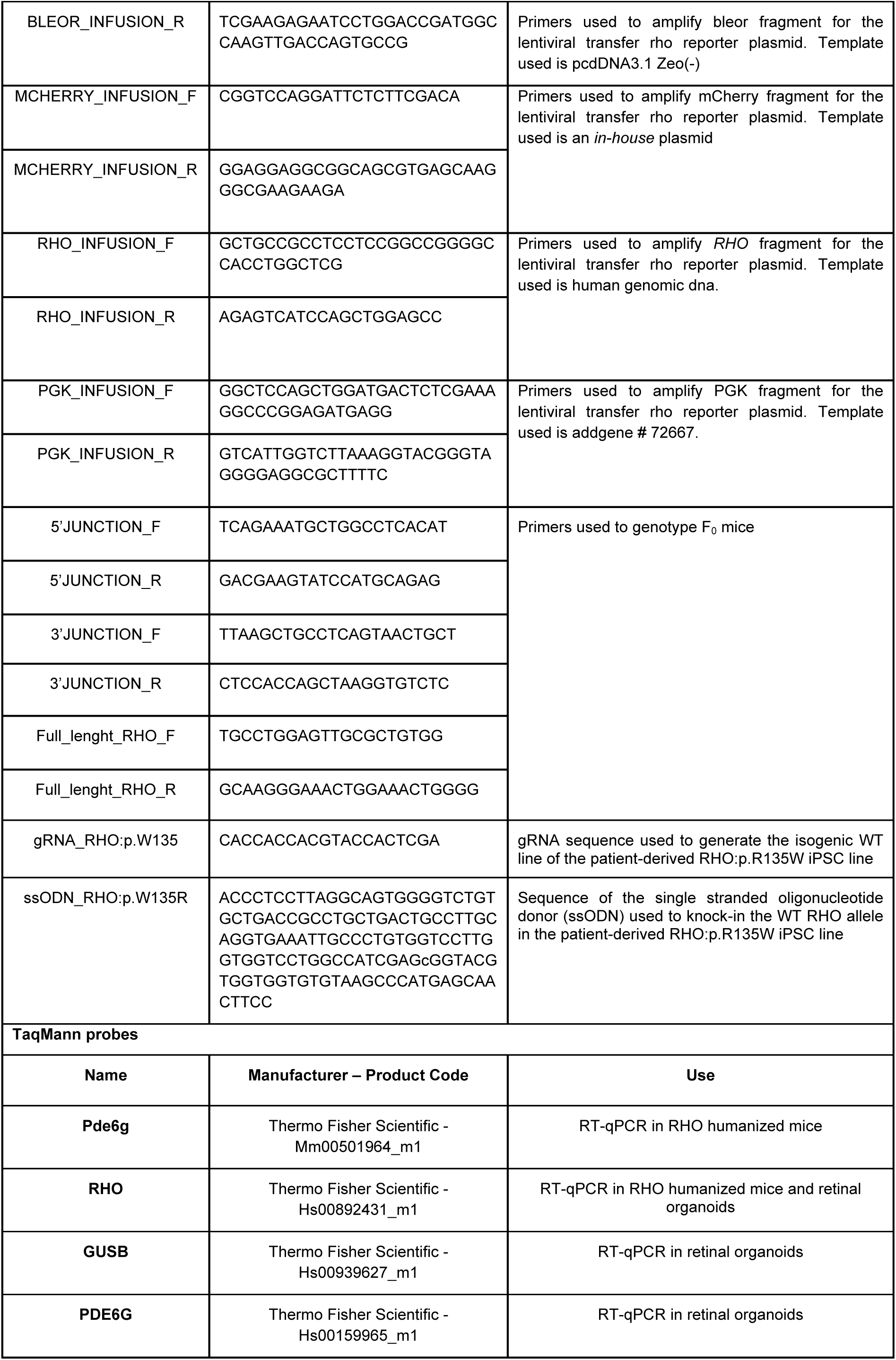
Primer list, gRNA, and ssODN and TaqMan probes

**Supplementary Table 3.** In-silico Off-target prediction File attached

**Supplementary Table 4.** Raw data mouse cDNA NGS

|  | Mouse # | Reads with A (%) | Reads with G (%) |
| --- | --- | --- | --- |
| Treated - LD | 1 | 34.41 | 63.62 |
|  | 2 | 33.08 | 64.73 |
|  | 3 | 36.41 | 61.66 |
|  | 4 | 31.93 | 65.58 |
| Counter - LD | 1 | 47.06 | 50.41 |
|  | 2 | 48.39 | 49.16 |
|  | 3 | 47.37 | 49.9 |
|  | 4 | 48.47 | 49.05 |
| Treated - HD | 1 | 30.03 | 67.95 |
|  | 2 | 29.70 | 68.19 |
|  | 3 | 29.85 | 68.35 |
|  | 4 | 26.12 | 71.89 |
| Counter - HD | 1 | 45.63 | 51.85 |
|  | 2 | 46.43 | 51.15 |
|  | 3 | 44.48 | 53.13 |
|  | 4 | 48.84 | 47.7 |

**Supplementary Table 5.**
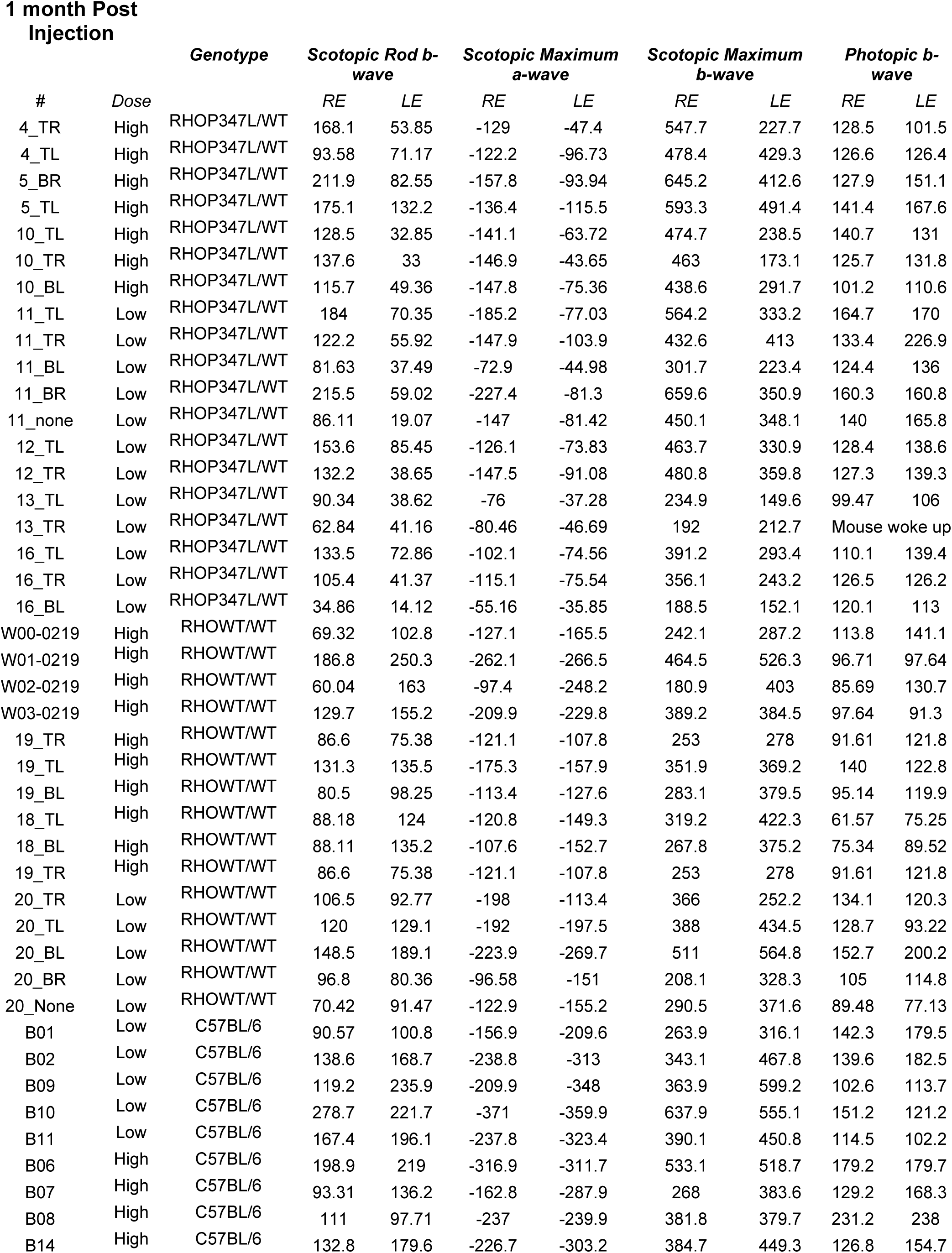
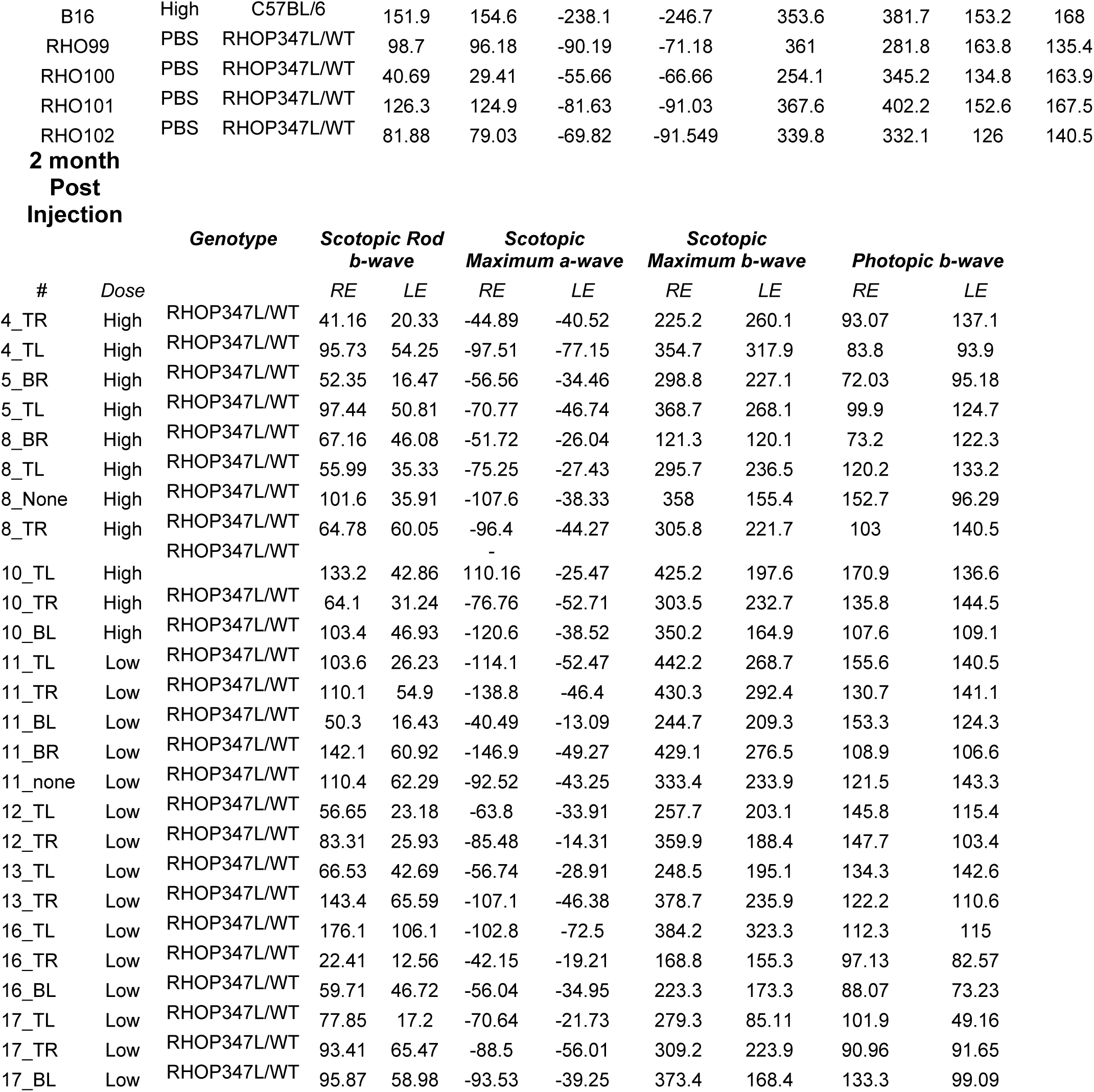
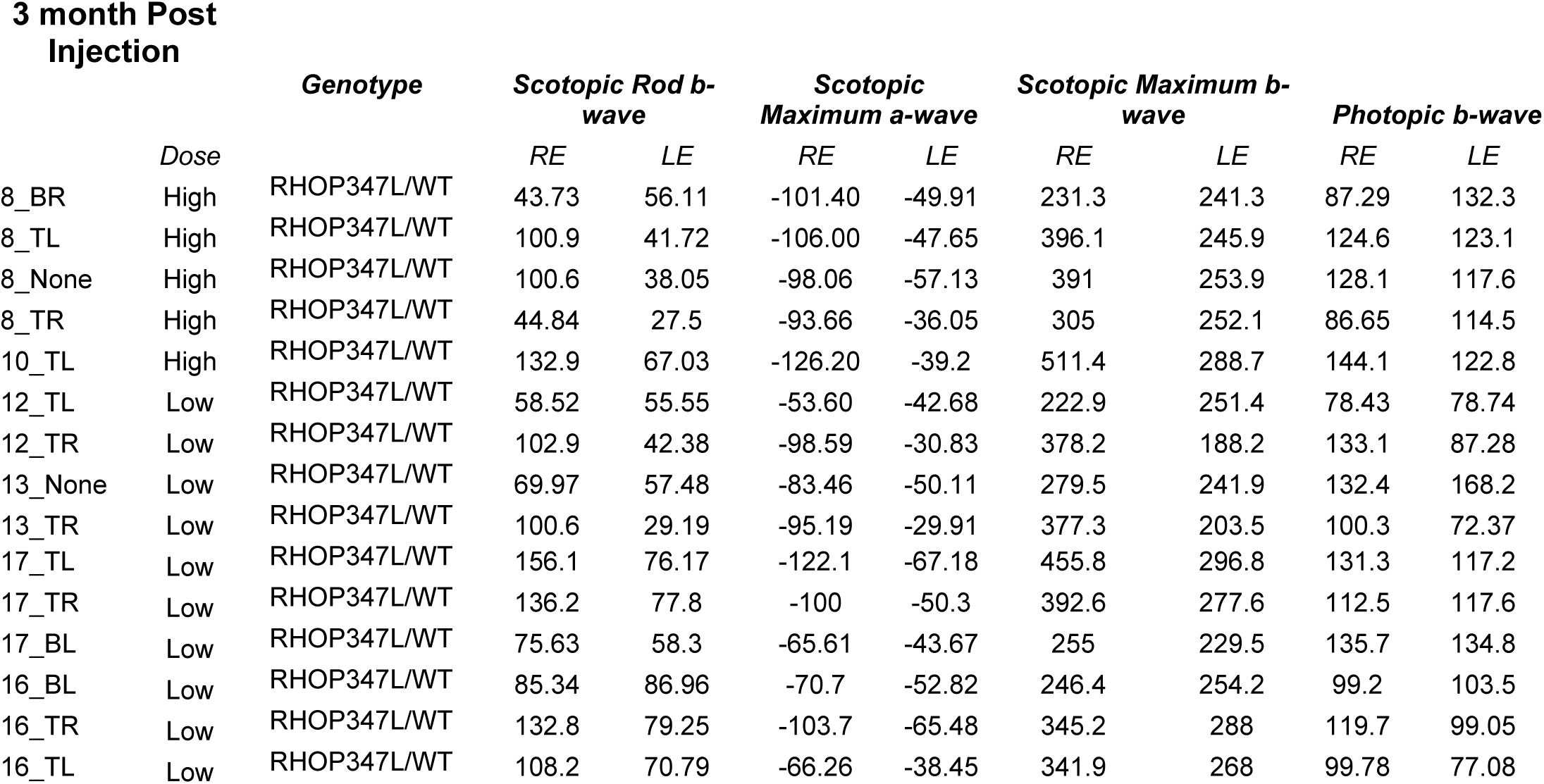
ERG responses

**Supplementary Table 6.**
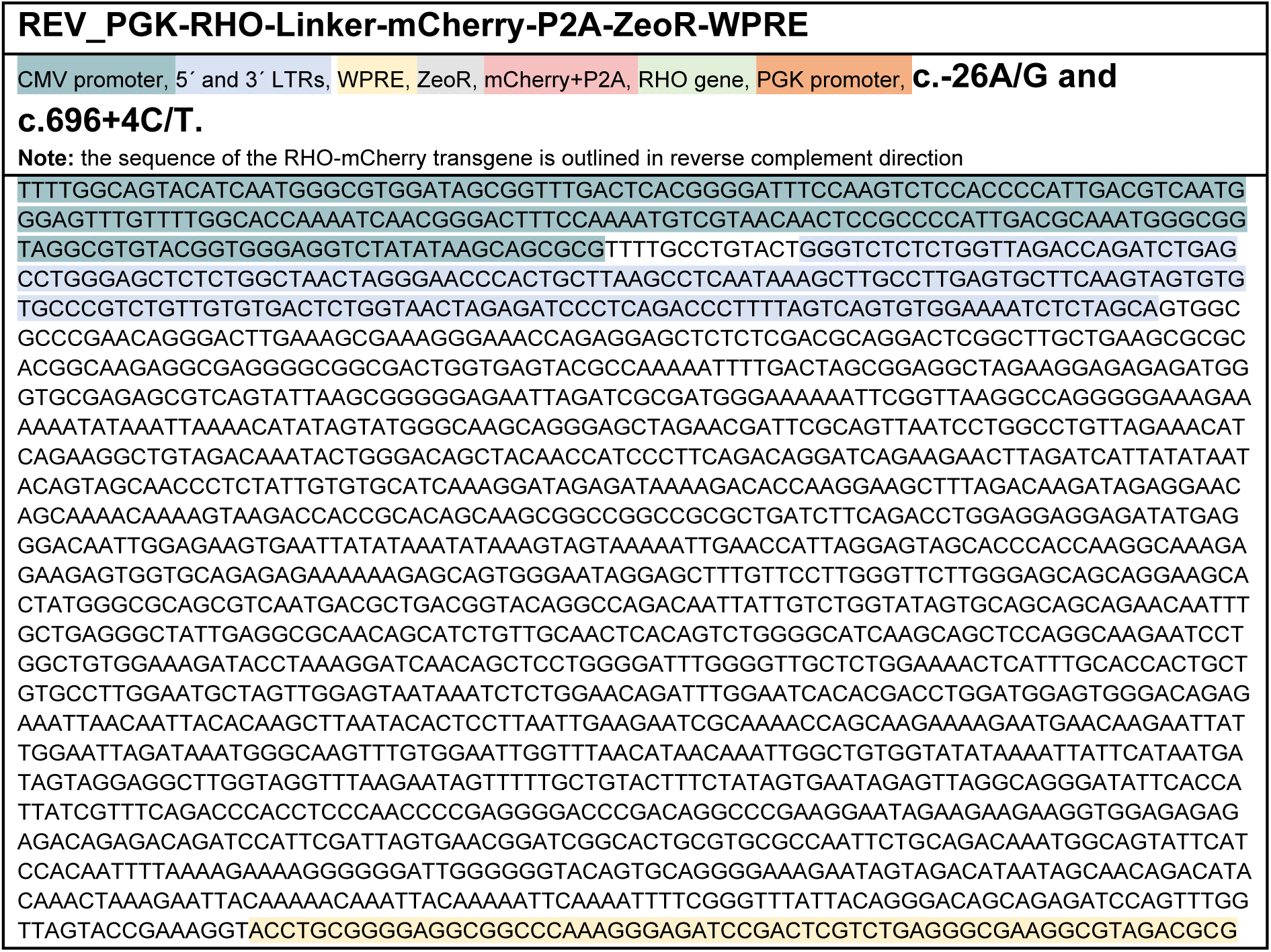
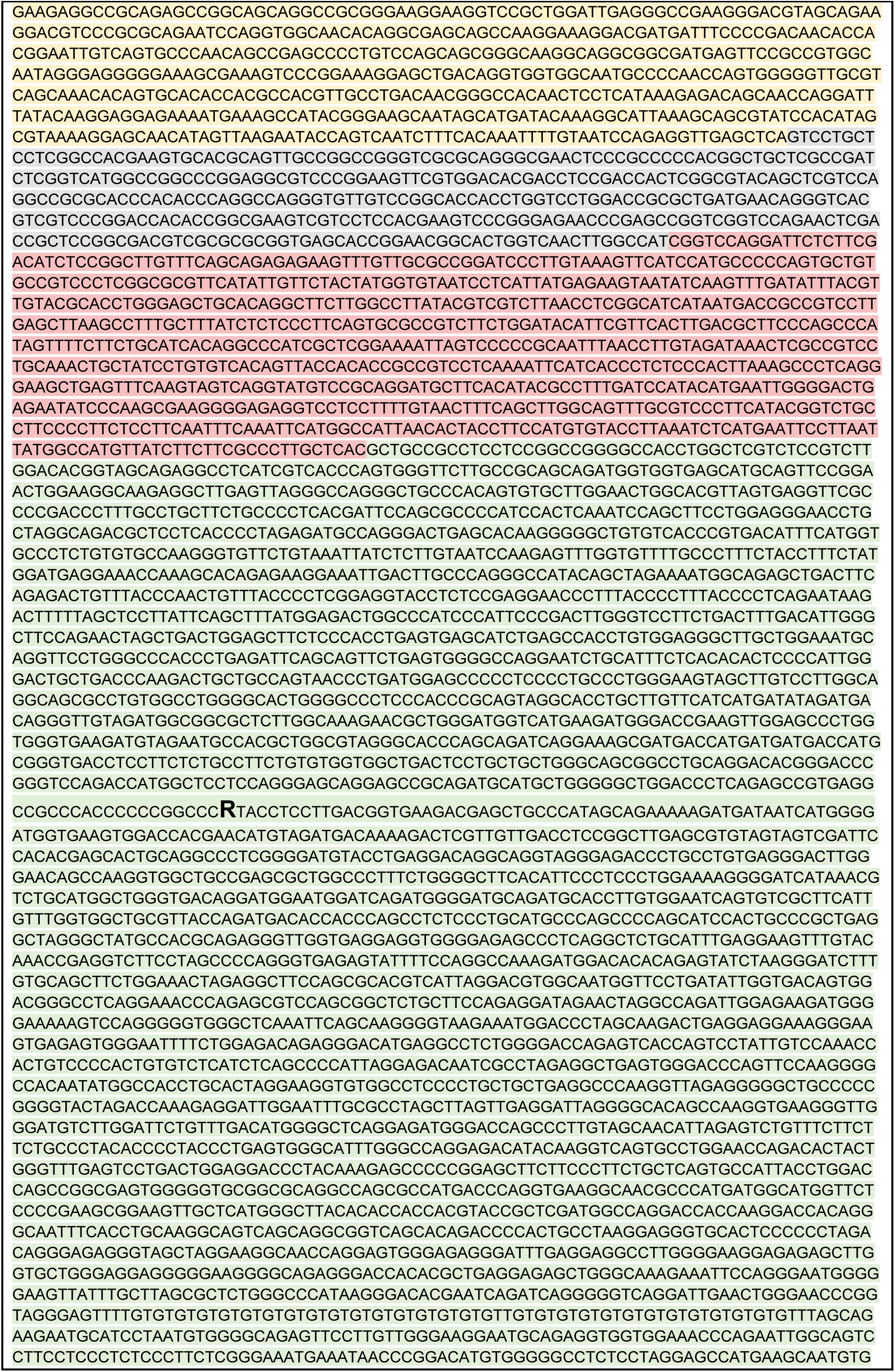
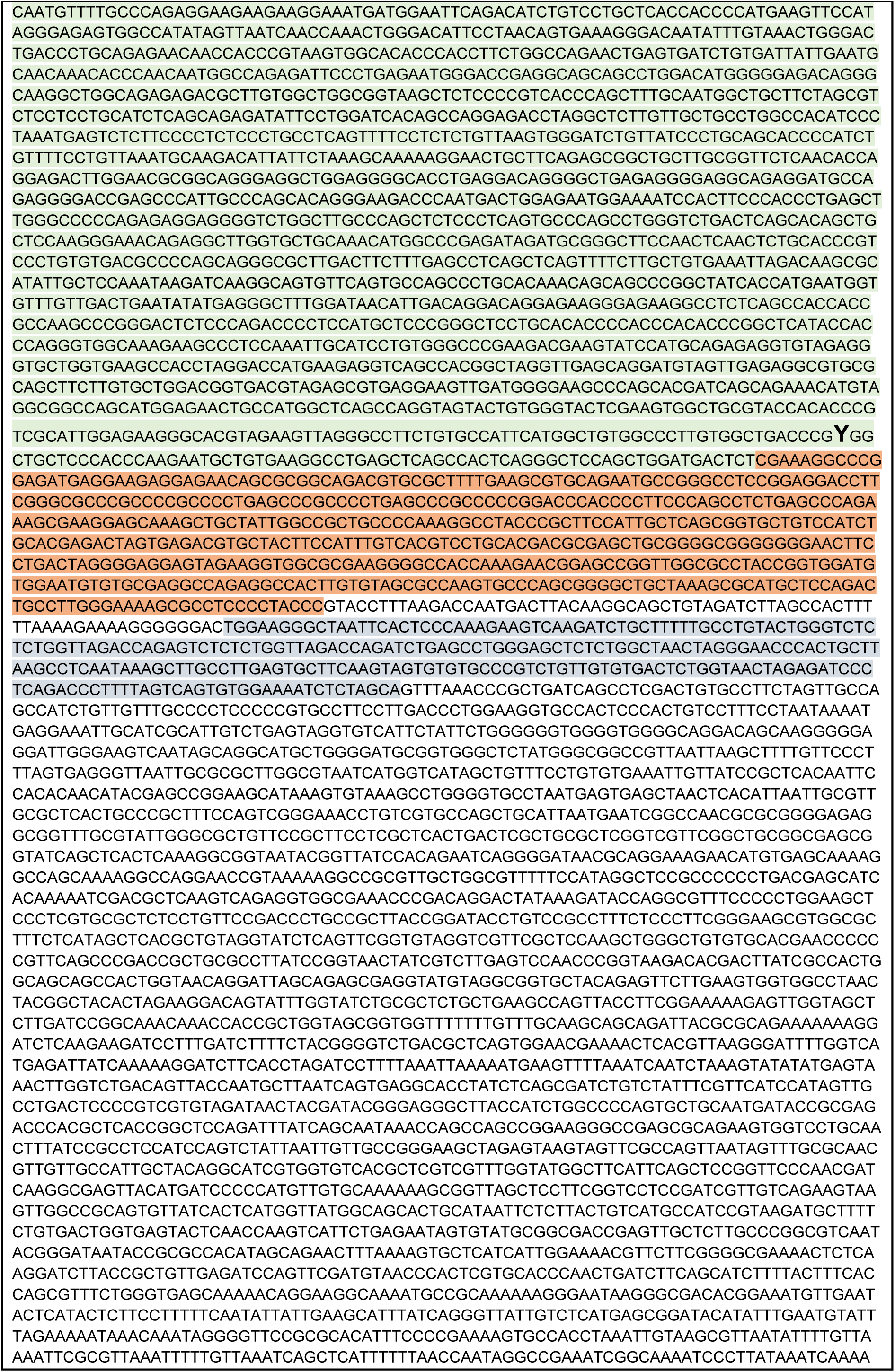
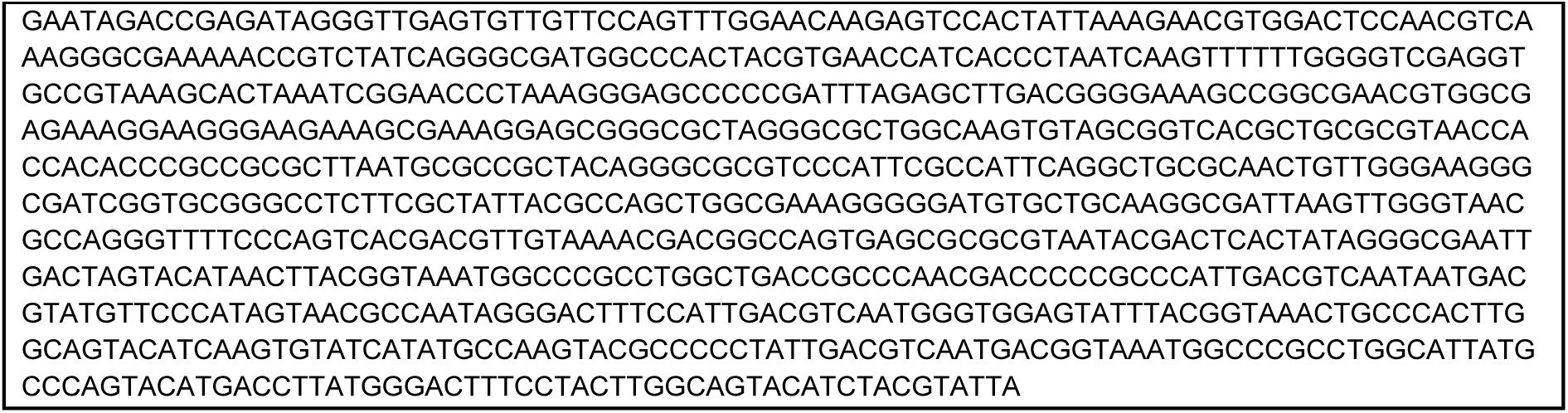
Plasmid sequences of REV_PKLV2.2-PGK-RHO-Linker- mCherry-P2A-ZeoR-WPRE. Lentiviral PKLV2.2-based transfer plasmids for generation of RHO-mCherry reporter cell lines. Plasmid for generation of cell lines *RHO*:c.[-26A;696+4T] and *RHO*:c.[-26G;696+4C] differ only in the two highlighted SNPs.

